# Hydrogen peroxide damage to scavenging function and ultrastructure of liver sinusoidal endothelial cells is prevented by n-acetyl-cysteine but not GSH

**DOI:** 10.1101/2024.08.26.609175

**Authors:** Larissa D. Kruse, Christopher Holte, Bartlomiej Zapotoczny, Eike C. Struck, Jasmin Schürstedt, Wolfgang Hübner, Thomas Huser, Karolina Szafranska

## Abstract

Reactive oxygen species (ROS) are prevalent in the liver during intoxication, infection, inflammation, and ageing. Changes in liver sinusoidal endothelial cells (LSECs) are associated with various liver diseases. We investigated how oxidative stress induced by H_2_O_2_ affects isolated rat LSECs at different concentrations (0.5-1000µM) and exposure times (10-120 min). Our findings show that H_2_O_2_ exposure affects several LSEC functions in a dose- and time-dependent manner: (1) cell viability, reducing potential, and scavenging function decreased as H_2_O_2_ concentration and exposure time increased; (2) intracellular ROS levels rose with higher H_2_O_2_ concentrations; (3) fenestrations exhibited a dynamic response, initially closing but partially reopening at H_2_O_2_ concentrations above 100µM after about 1 h; (4) scavenging function was affected after just 10 min of exposure, with the impact being irreversible and primarily affecting degradation rather than receptor-mediated uptake; (5) the tubulin network was disrupted in high H_2_O_2_ concentration while the actin cytoskeleton appears to remain largely intact. Finally, we found that reducing agents and thiol donors such as N-Acetyl Cysteine (NAC) and Glutathione (GSH) could protect cells from ROS-induced damage but could not reverse existing damage. Pretreatment with NAC, but not GSH, reduced the negative effects of ROS exposure suggesting that LSEC does not store an excess amount of GSH but rather can readily produce it in the occurrence of oxidative stress conditions. The observed thresholds in dose and time-dependent changes as well as the treatments with NAC/GSH confirm the existence of ROS depleting system in LSEC.

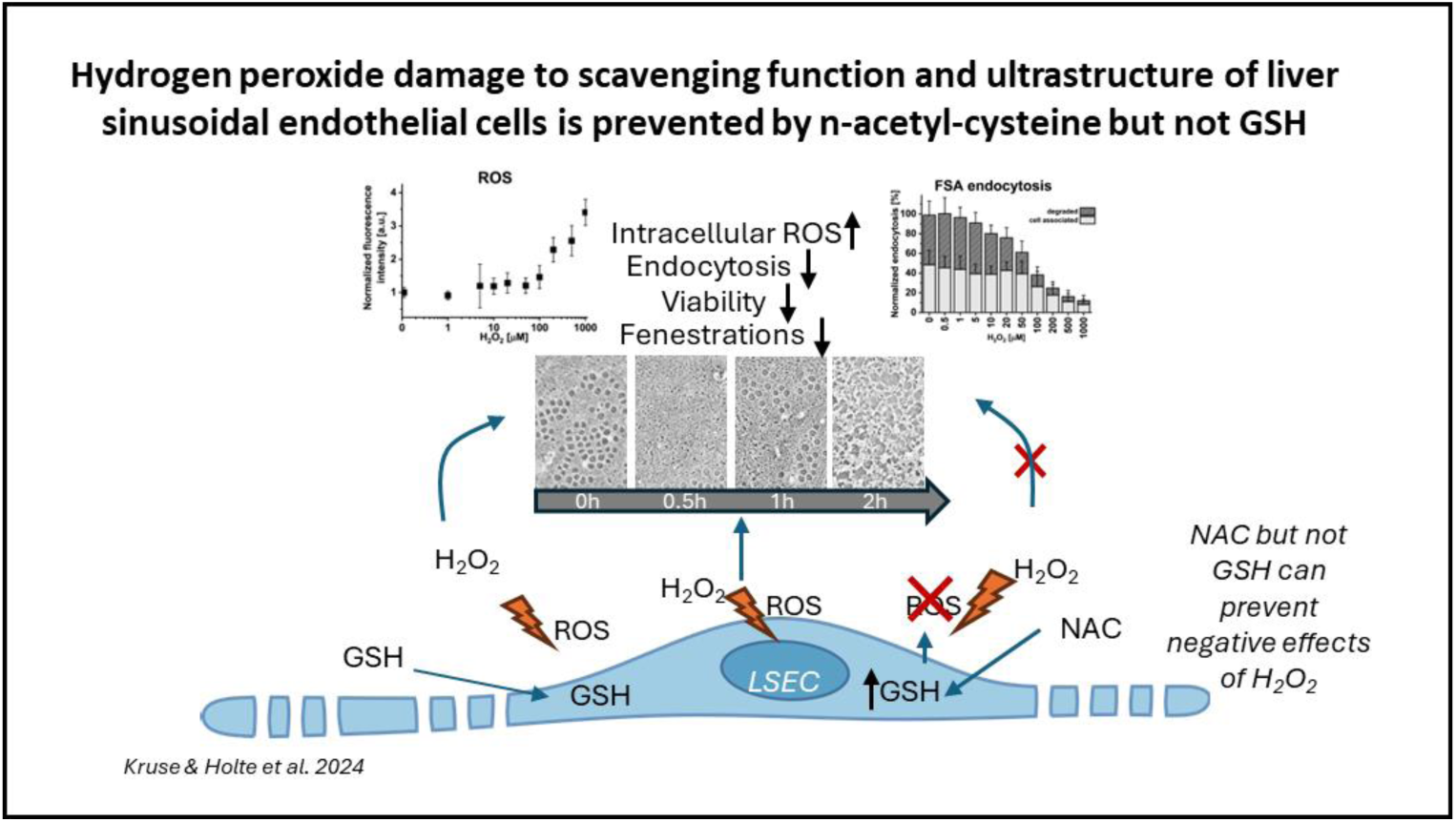

**Highlights:**

- ROS by H_2_O_2_ irreversibly depletes LSEC endocytic/scavenging function *in vitro*
- H_2_O_2_ exposure causes dynamic, dose-dependent defenestration of LSEC within 0.5 h
- Partial refenestration can occur after about 1h of exposure to H_2_O_2_
- NAC/GSH mitigate H_2_O_2_-induced ROS effects in LSEC
- LSEC do not store excess GSH but produce GSH when exposed to oxidative stress

## Introduction

Oxidative stress can be defined as an imbalance between the production, accumulation, and elimination of reactive oxygen species (ROS). ROS is a collective term for certain oxygen-containing oxidizing compounds, including but not limited to oxygen radicals. Radical ROS includes superoxide (O_2_^-•^), hydroxyl (HO^•^) and peroxyl radicals (HOO^•^) while nonradical ROS includes peroxide, hypochlorous acid, and ozone [1][2]. In biology, the role of ROS creates a paradox where the thin line between toxic and physiological effects is constantly being shifted. In homeostasis, ROS are present both intra- and extracellularly and can act as signaling molecules [3][4]. The mitochondria respiratory chain, lipoxygenases/cyclooxygenases, NO synthesis, nicotinamide adenine dinucleotide (NADPH) oxidases and xanthine oxidase are the main sources of intracellular ROS [5]. On the other hand, oxidative stress is associated with the pathogenesis of many diseases, ageing, promotion of inflammation and cytotoxicity [2][6]. The balance of the redox state in the cells is kept by a defensive system depending on enzymatic components, such as superoxide dismutase (SOD), catalase (CAT), and glutathione peroxidase (GPx), that protect the cells from ROS-induced cellular damage [7].

In the liver, ROS can be produced through endogenous (e.g. mitochondria, ER, peroxisomes) and exogenous (e.g. heavy metals, pollutants) sources [8]. Oxidative stress is implicated in the pathogenesis of diseases, drug-induced liver injury, reperfusion injury, sinusoidal obstruction syndrome, but also in inflammatory responses and ageing [6][9]. Even in non-pathological states, due to its location in the cardiovascular system, the liver is constantly the site of oxidative stress with portal blood and highly metabolically active hepatocytes being the main ROS sources. The portal vein is the main blood supply of the liver and the source of gut-derived toxins, and any other substance absorbed via the gastrointestinal tract. Cytochromes P450 enzymes present in hepatocytes are known to generate ROS in the liver [10]. Between these two sources of ROS lie the liver sinusoidal endothelial cells (LSEC) creating a barrier and regulating the bidirectional transport between blood and hepatocytes through the space of Disse [11][12]. This placement makes LSEC especially exposed to oxidative stress also from other liver sinusoidal cells such as liver resident macrophages – Kupffer cells, or stellate cells can be a source of ROS if responding to inflammatory stimuli [13][14].

Since LSEC are especially vulnerable to ROS stimuli, a defensive glutathione (GSH)-based system was proposed that allows maintenance of the redox balance under physiological conditions [15][16]. To deepen the knowledge about the LSEC and ROS interaction we investigated the time- and dose-dependent effects of H_2_O_2_ on rat LSEC morphology and functions *in vitro*. Moreover, by using the pre- and co-treatments with ROS depleting agents – NAC and GSH, we further study the anti-ROS defense mechanisms in LSEC.

## Materials and methods

### LSEC isolation and cell culture

The experiments followed protocols approved by the local Animal Care and National Animal Research Authority at the Norwegian Food Safety Authority (Mattilsynet; Approval IDs: 4001, 8455, and 0817). All experiments were performed in accordance with relevant approved guidelines and regulations, and reported following ARRIVE guidelines. All experiments were performed in a minimum of 3 bio replicates with the total use of research animals of 18 rats. Sprague Dawley male rats (Janvier, France; Strain: RjHan:SD) were kept under standard conditions and fed standard chow ad libitum.

Rats (body weight 300–500 g, age 2–3 months) were anesthetized with a mixture of xylazine and ketamine and LSECs were isolated using the modified protocol described in [17]. Briefly, after liver perfusion and digestion using Liberase^TM^ (Roche), parenchymal cells were removed by differential centrifugation, and LSEC were separated using primary mouse SE-1 antibodies (Novus Biologicals) and secondary immunomagnetic beads conjugated with anti-mouse IgG2a+b (MACS, MiltenyiBiotec). After isolation, cells were seeded according to the experiment type on surfaces coated with 0.2 µg/mL of human fibronectin for 10 min. Samples were incubated for 3 h in RPMI-1640 media (Sigma-Aldrich) at 37°C with 5% CO_2_/5% O_2_ before treatments.

### Treatments

H_2_O_2_ (hydrogen peroxide, VWR International) was selected as a ROS-inducing agents and n-acetyl cysteine (NAC, Sigma-Aldrich) and glutathione (GSH, Sigma-Aldrich) were used as ROS-depleting agents. Hydrogen peroxide and GSH solutions were freshly prepared for each experiment just before use. Stock solutions of NAC (50 mg/ml in water, pH 7.0) were prepared and stored in −20°C until use. All treatments were made in RPMI 1640 cell culture media.

For experiments involving pretreatments, samples were treated with GSH (500 µM)/NAC (0.5 mg/ml, 2 mg/ml) for 30 min and rinsed with prewarmed RPMI media before further treatment. The indicated treatment times do not include pretreatment.

### Viability assays

#### LDH

LSEC were seeded on standard 48-wellplates (300 000 cells/well) and commercial luminescence LDH detection kit (LDH-Glo^TM^, Promega) was used following the manufacturer’s protocol to assess cell viability. After treatments, 50 µl of media samples were collected at selected timepoints (0.5 h, 1 h, 2 h and 5 h) into 450 µl of freezing buffer (details in the manufacturer’s protocol) and stored in −20°C until measurements.

#### Resazurin

LSEC were seeded on standard 48-wellplates (300 000 cells/well) and a commercial Resazurin/resorufin assay (Biotechne) was used as an indicator of the mitochondrial function and viability. Together with the treatments 1:10 resazurin reagent was added to the culture media. After set timepoints (1 h, 2 h, 3 h, 4/5 h) the supernatants (50 µl) were collected and fluorescence measured using a plate reader (excitation 530-570 nm emission 580-590 nm).

### Scavenging assay

A radiolabeled formaldehyde-treated serum albumin (FSA)-based scavenging assay was used for quantitative studies of endocytosis and degradation in LSEC [18][19]. Fully confluent cultures of rat LSEC were established in 48-well culture dishes (300 000 cells/well). For experiments that included pretreatments, samples were treated with NAC/GSH for 30 min before the scavenging assay. Otherwise, directly after treatment, radiolabeled formaldehyde-treated serum albumin (^125^I-FSA) was added to each well (approximately 30 ng) together with human serum albumin (HSA) (final HSA concentration of 1%) and incubated for 2 h. Thereafter, the cell-associated and degraded FSA fractions were analyzed as described previously [18][19].

Alternatively, FSA fluorescently labelled with AlexaFluor488 was used for the qualitative assessment of endocytosis in LSEC populations. FSA was added to the wells for the last 15 min of the 2 h treatments with H_2_O_2_ cells were then fixed with 4% formaldehyde in PBS for 15 min and stored in PBS until imaging.

### Imaging

#### SEM

The detailed method of sample preparation and imaging for SEM was described in [20]. In this study, rat LSEC were seeded on fibronectin-coated 16-wellplates (CS16-CultureWell™ Removable Chambered Coverglass, Grace Bio-labs) with a density of about 60 000 cells/well (0.4 cm^2^) and after treatments fixed using McDowell’s solution (4% formaldehyde and 1% glutaraldehyde in PHEM buffer pH 7.2) [21] and stored in this fixative until further processing. Samples were then washed in PHEM before 1 h incubation with 1% tannic acid (freshly prepared in PHEM) followed by 1 h in 1% osmium tetroxide in H_2_O and dehydration in an ethanol gradient (30%, 60%, 90%, 4×100% for 5 min each) and chemical drying in hexamethyldisilazane twice for 2 min (Sigma-Aldrich, Oslo, Norway). Before imaging samples were mounted on aluminum stubs and coated with 10 nm layer of gold/palladium alloys.

#### SIM/fluorescence microscopy

Samples were permeabilized with 0.5% Triton X100 for 90-120 s and stained with anti-α-tubulin antibodies conjugated directly with AlexaFluor647 (Santa Cruz, Cat. Nr. sc-23948), at a concentration of 2 µg/mL in 1% PBST (PBS containing 0.05% Tween-20), overnight at room temperature (RT). After multiple washes with PHEM, 5 min, 15 min, 30 min and 1 h, the samples were incubated with Phalloidin-AlexaFluor555 (ThermoFisher, Cat. Nr. A34055) for 90 min at RT. Thereafter, samples were washed again for approximately 30 min and incubated with DAPI (Merck, Cat. Nr. D9542) for 10 min. After two PHEM washes and one wash with ddH_2_O, the samples were mounted on microscope slides with ProLong glass anti-fade mounting medium (ThermoFisher, Cat. Nr. P36982) and kept at 4°C in the dark until imaging with widefiled fluorescence microscope (EVOS M5000, ThermoFisher) or super-resolution structured illumination microscope The SIM images were acquired using a DeltaVision OMX 4.0 imaging system (GE Healthcare, Chicago, IL, USA) equipped with a 60X 1.42NA oil-immersion objective (Olympus, Tokyo, Japan) and four sCMOS cameras. We used 488, 568, and 642nm lasers for excitation. Image deconvolution and 3DSIM reconstructions were completed using the manufacturer-supplied softWoRx program (GE Healthcare). Representative 3D-SIM images were presented as Z-projections."

#### AFM

LSEC were isolated and cryopreserved as described previously in [22]. After thawing, cells were seeded on the fibronectin-coated Petri dish (TPP) and kept in the incubator (37 °C and 5% CO_2_) for 4-8 h in the EGM-2 (Lonza) containing 2% FBS. Prior to measurements, the cell medium was replaced with fresh medium containing 25 mM HEPES and placed on the AFM microscope. All measurements were conducted using Nanowizard 4 (JPK Instruments) atomic force microscope. The microscope was equipped with a temperature controlling system, PetriDish heater (JPK Instruments). The experiments were performed in 37°C. Cells were measured with silicon nitride cantilevers (SCM-PIC-V2, Bruker) characterized by a nominal spring constant of 0.1 N/m and a nominal tip radius of 25 nm. All measurements of cell dynamics in response to hydrogen peroxide were performed using Quantitative Imaging™ mode according to the methodology described before [23][24]. Briefly, in each pixel point of the image, an independent force– distance (FD) curve was performed. A set of FD-curves was used then to reconstruct images of cell topography for selected loading force and images representing stiffness distribution.

The loading force used for QI measurements ranged from 0.2 to 0.3 nN and was adjusted to the scanning conditions for individual cantilevers. Firstly, LSEC was selected using an optical microscope and a low-resolution AFM image was collected. The area of interest, covering several sieve plates, was then selected in order to obtain a time per frame in range of minutes and a pixel point resolution that allows for easily distinction of individual fenestrations. After collecting few frames, H_2_O_2_ was injected into the culture medium and the same area was repetitively scanned. Several images collected consistently in the same area were presented in the form of supplementary videos allowing to track cell dynamics and changes in fenestration number with time. The experiments were conducted in 3 independent experiments.

### ROS detection assay

A commercially available ROS indicator was used for detection of intracellular ROS (CM-H2DCFDA, Invitrogen, USA) according to the manufacturer’s protocol. LSEC were seeded on standard 48-wellplates (250 000 cells/well) and after about 2 hours incubation the cell media was exchanged to Hank’s buffered salt solution (HBSS, with Ca^2+^/Mg^2+^, glucose, no phenol) containing 3 µg/ml of the dye and incubated for 45 min in 37°C (without CO_2_). Afterwards, cells were rinsed and incubated for additional 20 min in RPMI (loading time) before treatment with selected agents. Fluorescent images were taken from live cells in HBSS directly after 60min treatments and analyzed using ImageJ/Fiji [25] to compare fluorescence intensity signal.

### Image analysis and statistics

SEM images were analyzed using semi-automatic methods as described in [20]. Fenestration size, porosity and frequency were assessed in each sample from randomly selected areas where 15-20 cells were imaged. Open pores with diameters between 50 and 350 nm were defined as fenestrations and holes larger than 350 nm as gaps. The mean fenestration diameter was calculated as a peak of the Gaussian fit in the histogram combining data from all measured fenestration from each treatment group. Porosity was defined as the sum area of fenestrations per total area of the cell in the micrographs. Frequency was denoted as the total number of fenestrations per sum area of the cell excluding the sum area of gaps.

Statistical analyses were performed using R statistical software [26]. Data normalization and transformation steps were conducted as per experimental requirements to ensure comparability across conditions. The Shapiro-Wilk test was employed to assess the normality of the data distribution within each group. Based on the normality test outcomes, data were subjected to either parametric or non-parametric statistical analyses. Significance analysis was performed on non-normalised data.

For time-dependent data a model analysis (as p for trend) was performed to evaluate the effects of treatments, and, where relevant, their interaction with time points on the measured outcome, separately for each time point. For normally distributed datasets with homogeneity of variances, Analysis of Variance (ANOVA) was performed to evaluate the main effects of treatments and, where applicable, their interaction with time points on the measured outcome. This included One-Way ANOVA in the absence of time-dependent variables to assess the impact of different treatments on the outcome variable, and Two-Way ANOVA with Interaction used for time-dependent datasets to not only assess the main effects of treatment and time and potential interaction between these two factors. Post-hoc analyses, utilizing Tukey’s Honestly Significant Difference test, were conducted following significant ANOVA results to identify specific pairwise differences between treatment groups.

In cases where data failed to meet normality criteria, the Kruskal-Wallis test was used to discern statistical differences across treatment groups. For time-dependent datasets, this analysis was executed individually for each time point. Subsequent post-hoc Dunn tests, with Benjamini-Hochberg correction for multiple comparisons, were used to analyse group specific differences.

Analysis outcomes, including fold changes, estimates of central tendency, variance, and statistical significance levels (p values) can be found in supplementary table X. Significance was denoted using conventional asterisks notation to indicate varying levels of statistical significance (p >= 0.05 = ns, p < 0.05 = *, p < 0.01 = **, p < 0.001 = ***). The processed data, alongside statistical summaries, were compiled into structured formats for visualization and further analysis.

## Results

### Effects of ROS on LSEC scavenging function and viability

First, a concentration range of the ROS-inducing factor – hydrogen peroxide – was tested to establish the amount necessary to change intracellular ROS levels in LSEC. A fluorescent-based ROS detection assay was used on LSEC challenged with increasing concentrations of hydrogen (Figure 1A). No detectable ROS increase was observed for H_2_O_2_ concentrations below 5 µM and a small elevation of ROS was detected for medium concentrations (5-50 µM H_2_O_2_). A dose-dependent increase was observed for high (100-1000 µM) concentration of H_2_O_2_ reaching a 3.5-fold increase at 1000 µM.

**Figure 1.**
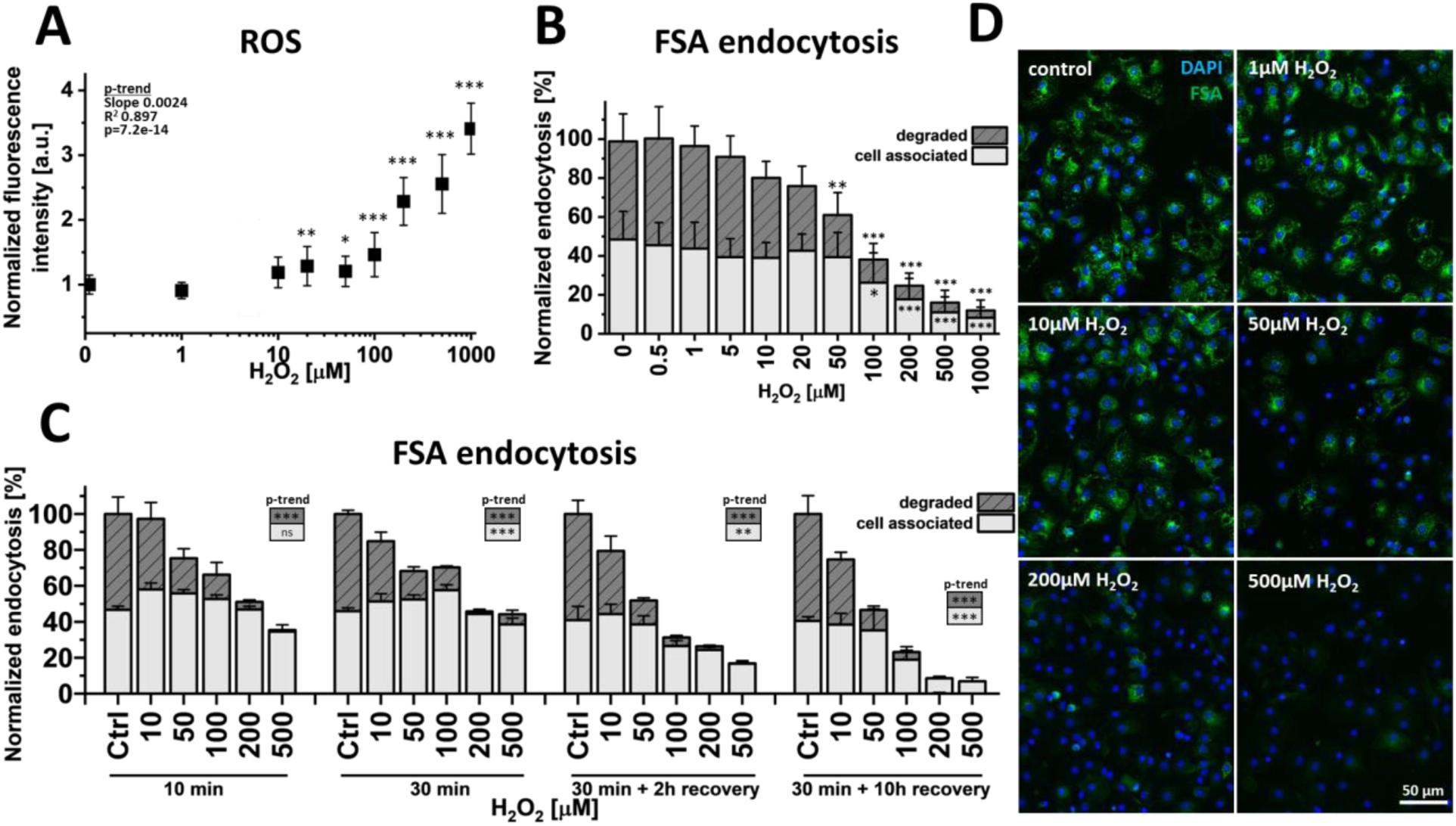
Effects of ROS on endocytosis of trace amount of formaldehyde-treated serum albumin (FSA) in rat liver sinusoidal endothelial cell (LSEC). (A) Intracellular ROS level after 1 h treatment with 0-1000 µM hydrogen peroxide. Fluorescence based assay data was normalized to the untreated control to show the fold-increase. Mean ±SD. (B,C) The effects of H_2_O_2_ on scavenging of ^125^I-FSA in LSEC. The total endocytic activity was 40-60% of the added radioactivity and was normalized to the untreated control. (B) Cells were treated with hydrogen peroxide together with ∼30 ng/ml of radiolabeled FSA for 2 h. (C) Cells were treated for 10/30 min with hydrogen peroxide then rinsed and either directly incubated with ∼30 ng/ml of radiolabeled FSA for 2 h or recovered with RPMI media for 2 h or 10 h before 2 h incubation with radiolabeled FSA. Dark/striped bars represent degraded fraction while light bars represent a cell-associated fraction of the ligand (±SD), (A/C) n=3-4 bio replicates, (B) n=7. (D) Uptake of FSA-AF488 after treatment with hydrogen peroxide shows a heterogeneous population of LSEC. FSA was added for the last 15 min of the 2 h treatment with H_2_O_2_. Significance indicated in relation to the control (A-B) or as p-for-trend analysis (A,C), ns – not significant, * p<0.05, ** p<0.01, *** p<0.001,details in Table S1-3.

The ROS effect on the scavenging system was studied using formaldehyde-treated serum albumin (FSA) based assays (Figure 1B-D). FSA is a ligand for stabilin 1 and 2 [27][28] and allows for assessment of both uptake and degradation by the scavenging system in LSEC [19]. Low concentrations of hydrogen peroxide (below 5 µM) did not affect scavenging, while in medium concentrations of 5-50 µM a steady decrease in the degraded fraction of FSA was observed, up to a 40% reduction. For high concentrations of hydrogen peroxide (>100 µM) a significant drop in both degraded and cell associated fraction of FSA occurred with almost complete inhibition of degradation at 1000 µM H_2_O_2_ (see Table S2 in supplemental for detailed statistical analysis). A p for trend analysis revealed a significant shift downwards for the degraded (p_degraded_ = 1.00E-09 (***)) and cell associated (p_cell_associated_ = 2.57E-08 (***)) fractions over increased doses of H_2_O_2_. We applied a split model to determine the effect threshold. P for trend analysis for degraded and cell associated values was split into doses from 0-50 µM H_2_O_2_ and 50-1000 µM. The latter showed a significant reduction for both models (p_degraded_50-1000_ = 0.009, (**), p_cell_associated_50-1000_ = 0.0005 (***)), while 0-50 µM H_2_O_2_ was significant only in the degraded fraction (p_degraded_0-50_ = 2.59E-06 (***)), showing a generally stronger effect at the higher concentrations of H_2_O_2_ with more influence on the degradation than uptake (statistical results can be seen in Table S1). The strength of the effect on scavenging in low/medium/high concentration ranges of hydrogen peroxide greatly resembles the detected intracellular ROS levels. Similar trends were observed with a scavenging assay based on radiolabeled collagen α-chain – a specific ligand of mannose receptor (Supplementary information in Figure S1).

These results were confirmed in qualitative experiments using fluorescently labelled FSA where endocytosis ligands were added for the last 15 min of the 2 h treatment with H_2_O_2_ (Figure 1D). No FSA uptake was observed for concentrations above 500 µM, while for concentrations of 50-200 µM only a fraction of the LSEC population showed remaining endocytic activity. This finding suggests that the decrease in the scavenging function is a result of the decreasing number of cells that can efficiently perform endocytosis rather than lower endocytic activity per cell.

To better understand the dynamics and reversibility of the H_2_O_2_ effect, different durations of ROS induction in LSEC were studied using the quantitative/radiolabeled scavenging assay (Figure 1C). Firstly, LSEC were treated with different doses of H_2_O_2_ for 10 min and 30 min before the addition of FSA. Both treatments of 10 min and 30 min showed a decrease in the endocytic activity for H_2_O_2_ concentrations above 10 µM. Additionally, we investigated whether the scavenging systems remained irreversibly damaged. To test this, recovery times of either 2 h or 10 h were applied after initial the 30 min challange with H_2_O_2_ in a range of concentrations. The results suggest that scavenging systems remain irreversibly damaged despite the recovery time and a further decrease of FSA uptake was observed 10 h after the initial treatment. The p for trend analysis shows a significant reduction in degraded fractions over all experiments in Figure 1C. The cell associated fraction shows a significant shift for 30 min with no recovery time albeit with a positive slope, suggesting it is driven by the upwards trend between 0 and 100 µM of of H_2_O_2_ (see Table S3 for all values).

The changes in measured scavenging activity can be a result of either a disruption of the scavenging system or damage to the cell resulting in cell death. In the next step, following viability assays were used to verify this.

Two approaches were applied to investigate the effects of hydrogen peroxide inducing ROS on LSEC viability. Functional viability was assessed using the resazurin assay (Figure 2A), while structural integrity was studied using the lactase dehydrogenase (LDH) release viability assay (Figure 2B).

**Figure 2.**
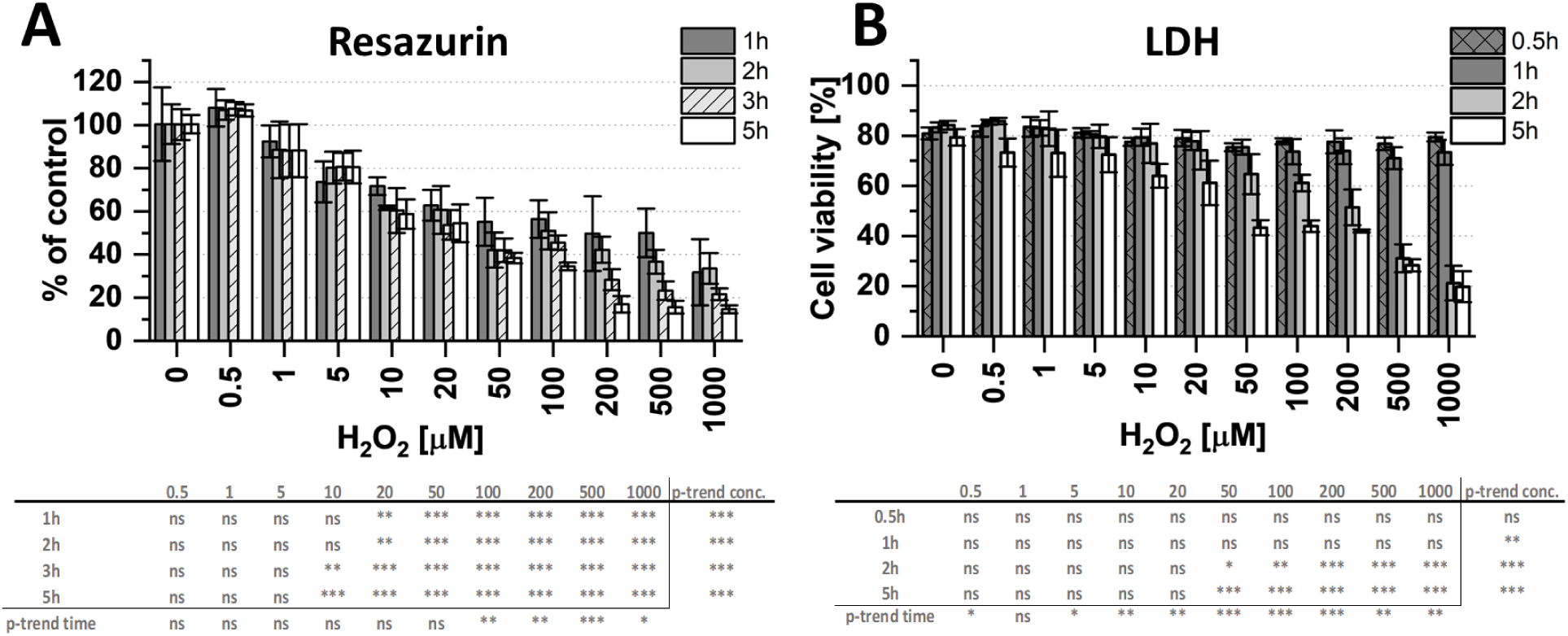
Influence of hydrogen peroxide induced ROS on rat LSEC viability. Functional viability/reducing potential of the cell was assessed using resazurin/resorufin assay (A) and structural viability/cell integrity was studied with LDH-release assay (B). Measurements were conducted under continuous treatment with hydrogen peroxide on selected timepoints (0.5-5h). The cell viability was calculated and normalized to the untreated control for resazurin assay and to the total LDH after cell lysis for LDH-release assay. Average data from 3 independent experiments/bio replicates ±SD is presented. Statistical analyses below graphs show p-values for trend analysis in time and concentration dependency. ANOVA for different time and concentrations compared to control values of non treated cells.

Resazurin assay measurements correspond with the reducing power of the cell often connected to mitochondrial metabolic activity. A significant dose-dependent decrease was observed for all timepoints for concentrations in the range of 20-1000 µM, while a time-dependent decrease occurred for concentrations of 100 µM H_2_O_2_ and higher. For treatments in the range of 0.5 µM to 100 µM H_2_O_2,_ the initial decrease observed after 1 h of treatment did not progress in further time points. For concentrations of 100 µM H_2_O_2_ and higher., the initial effect after 1 h was followed by further progressive reduction until reaching below 20% of the control after 5h of treatment.

Upon cell membrane damage, LDH is released into the culture medium where it can be quantitatively measured; as it describes the membrane integrity, we define it as a structural viability. The initial structural viability of 80% in untreated cells is related to cell death occurring during the first hours after isolation in primary LSEC. The treatment with hydrogen peroxide did not affect the structural viability in the first 1 h for any of the concentrations. However, after 2 h and 5 h of treatment with H_2_O_2_ a significant dose-dependent increase in LDH release in comparison with the control was observed for treatments with 50 µM and above.

Both the Resazurin and LDH release assay results suggest that although for the H_2_O_2_ concentrations below 50 µM the cells are affected rapidly after the exposure but afterwards remain stable without further damage. On the other hand, in high concentrations of hydrogen peroxide, above 50 µM, the functional and structural viability progressively decreased in time. This pattern points out the existence of a ROS-depleting system in LSEC that can mitigate the negative effects of ROS until a certain concentration threshold.

### ROS-induced changes in LSEC morphology

Table S5). Additionally, distorted sieve plates resembling previously reported defenestration centres (DFC)[29][24] were observed for H_2_O_2_ concentrations above 5 µM (Figure 3B,C). After the first 0.5h of treatment near complete defenestration was observed for 100-500 µM H_2_O_2_ while after 1 h of treatment, the number of fenestrations increased, however never returning to the control levels (Figure 3B,H). Significant, biologically relevant differences in fenestration frequency could be observed between the control and 500 µM H_2_O_2_, for 1 h treatment and between 30 min and 1 h treatment of 500 µM H_2_O_2_ (see ^T^able S^4^ for p-values). The detailed fenestration frequency data shows that the cell population became heterogeneous with some cells remaining defenestrated while others regained the porous morphology (Figure 3I). For high (>100 µM) H_2_O_2_ concentrations nearly no viable cells were observed after 2 h, with a majority of the sample presenting distorted/discontinued cell membranes suggesting necrotic cells death without typical shrunken apoptotic morphology (Figure 3D). This observation confirms the previous structural viability data where significant increase in the LDH release was detected only after 2h of treatment with >100 µM H_2_O_2_ with no significant increase for 0.5h and 1h(Figure 2B). In samples treated with H_2_O_2_ concentrations below 20 µM a dose-dependent loss of fenestrations was observed for all selected time points.

**Figure 3.**
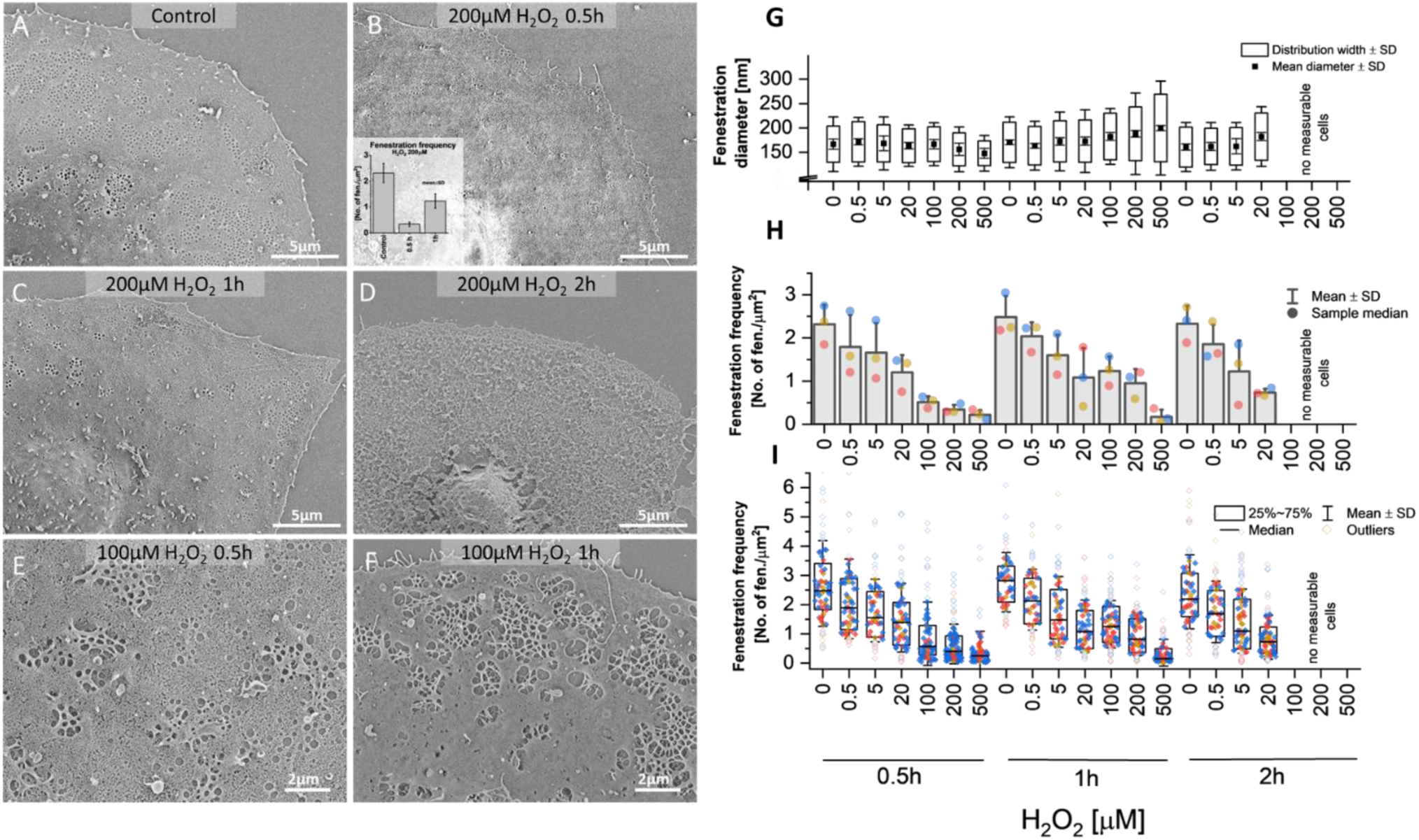
Changes in LSEC fenestrated morphology after exposure to H_2_O_2._ (A-F) Representative scanning electron microscopy images and (G-I) quantitative analysis results of hydrogen peroxide treated rat LSEC. A clear dose-dependent effect was observed for all timepoints (0.5-2h). Treatment with 100-200μ M H_2_O_2_ resulted in an initial decrease in fenestration number after 0.5h followed by an increase in fenestration number after 1h in a part of cell population and later degradation of cell membrane after 2h of treatment. (G-I) SEM images of rat LSEC treated with 0-500 µM H_2_O_2_ for 0.5-2h were analyzed to obtain parameters such as fenestration diameter, fenestration frequency and porosity (Figure S2)(G) Mean±SD was calculated from medians of fenestration diameters for each sample and distribution widths were calculated at the half maximum of the Gaussian distribution fit. (H) Each dot represents the mean fenestration frequency value calculated from each bioreplicate (all datapoints shown in (I)). (I) Each point represents data from a single cell and each color an individual bioraplicate. The effects of ROS on LSEC morphology were studied using electron and light microscopy. The detailed morphological structure was observed with scanning electron microscopy (Figure 3A-F) and images were quantitatively analyzed to calculate fenestration diameter, fenestration frequency and porosity (Figure 3G-I, Figure S2). For all time points, a significant, dose-dependent reduction in the number of fenestrations was observed (p for trend analyses: p_0.5h_ = 0.0002 (***); p_1h_ = 4.59E-05 (***); p_2h_ = 0.0013 (**),

The fenestration diameter in samples treated with high H_2_O_2_ concentrations above 100 µM showed a dose-dependent decrease. In particular, for 500 µM H_2_O_2_ treatment fenestration diameter decreased from 167 nm to 147 nm after 0.5 h, and later increased to 200 nm after 1 h (Figure 3G). A similar trend was observed for fenestration diameter distribution width which initially decreased after 0.5 h and then increased after 1 h for concentrations of 200-500 µM H_2_O_2_. In similar range of concentrations of 100-1000 µM, after 2 h of treatment it was not possible to distinguish fenestrations in SEM images due to disturbed cell membranes (Figure 3D).

The combination of initial defenestration followed by reopening of fenestrations and both time- and dose-dependent changes in the fenestration diameters suggest a dynamic response of LSEC morphology to hydrogen peroxide-induced ROS. To better understand this effect in temporal resolution we used Atomic Force Microscopy (AFM) for live imaging of LSEC treated with H_2_O_2_ (Figure 4).

**Figure 4.**
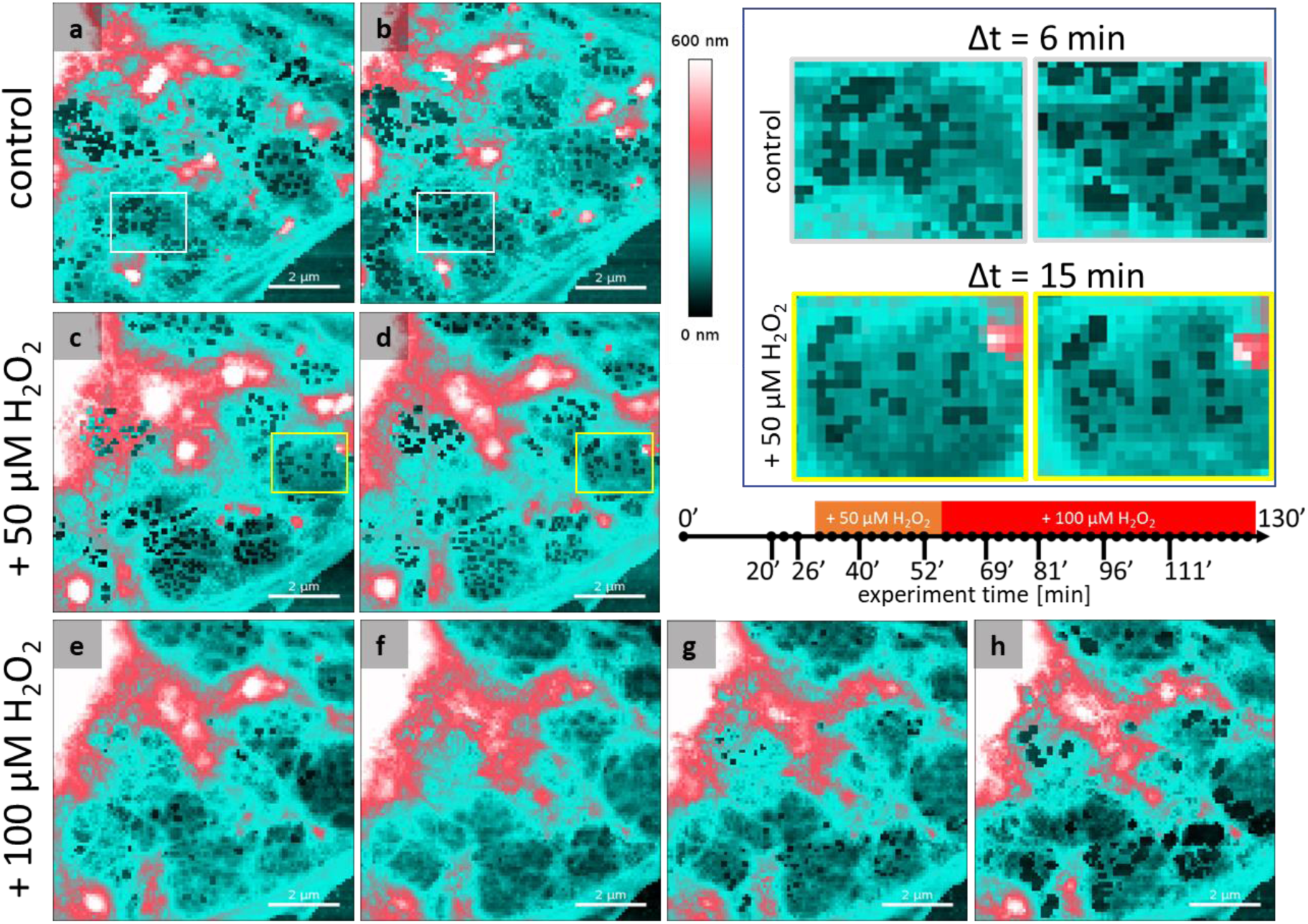
Dynamic defenestration and refenestration in live LSEC treated with hydrogen peroxide. Selected LSEC periphery with several sieve plates was scanned with AFM for 30 min revealing normal fenestration dynamics (white squares, a/b). After treatment with 50 µM H_2_O_2_ the processed of reduction of fenestration dynamics (yellow squares, c/d) and reduction of fenestration number were observed. The addition of H_2_O_2_ to a total concentration of 100 µM resulted in further gradual closing of all fenestrations in the following ∼20 l min and reopening in the following 30 min. Image size, resolution, and scanning speed were adjusted to obtain a final acquisition of 4 minutes per frame, each represented as a single point on the timeline. Image size and resolution: 8.5 × 8.5 μm, 100 × 100 pixels) (Scale bar = 2 μm). Presented images a-h corresponds with time points indicated on a timeline and were selected from 130 minutes long experiment, presented as Supplementary video SV1.

Highly dynamic fenestrations and sieve plates changing their size and position with time were detected at the beginning of the experiment. After treatment with 50 µM H_2_O_2_ we observed a reduction in fenestration number, confirming the SEM data from Figure 3. Moreover, the reduction of fenestration dynamics occurred nearly immediately after the injection of the agent into cell culture. The challenge with an additional 50 µM H_2_O_2_ (100 µM in total) resulted in the gradual closing of fenestrations within 20 min. Still, fenestration associated cytoskeleton ring (FACR) structure could be easily distinguished (Figure 4e/f), while the fenestrations remained closed and the cell membrane fused. During the following 30 min, we observed a gradual reopening of fenestrations. Reopened fenestrations quickly increased their dimensions often exceeding 300 nm. Newly formed fenestrations did not migrate within the cell and remained arrested in the same position. After an additional 30 min, we observed cell flattening at the peripheries with numerous fenestrations and gaps (Supplementary video X). Similar results were reproduced in two independent experiments, confirming the SEM results and providing a detailed understanding of the dynamics of this process.

### ROS-induces changes in LSEC cytoskeleton

In LSEC, both fenestrated morphology and scavenging functions are closely connected with the cytoskeleton. Therefore, we studied the changes in actin and tubulin under the influence of H_2_O_2_ using super-resolution optical nanoscopy to visualize the fine structure of the LSEC cytoskeleton. In high concentrations (>100µM), hydrogen peroxide disrupted the tubulin structure with numerous cells presenting a visibly reduced number of tubulin fibres (Figure 5). Moreover, in the affected cells microtubules seem to lose their characteristic organization – fibres emerging from the centrosome and surrounding sieve plates, and independent unconnected fibres were observed. For high concentrations of H_2_O_2,_ FSA was observed evenly scattered within the cell body and no longer followed the tubulin fibres suggesting disturbed transportation of the endocytic vesicles. No significant changes or stress fibre formation were observed in the actin cytoskeleton. Undisturbed actin mesh and regular fenestration associated cytoskeleton were observed in both treated and untreated cells.

**Figure 5.**
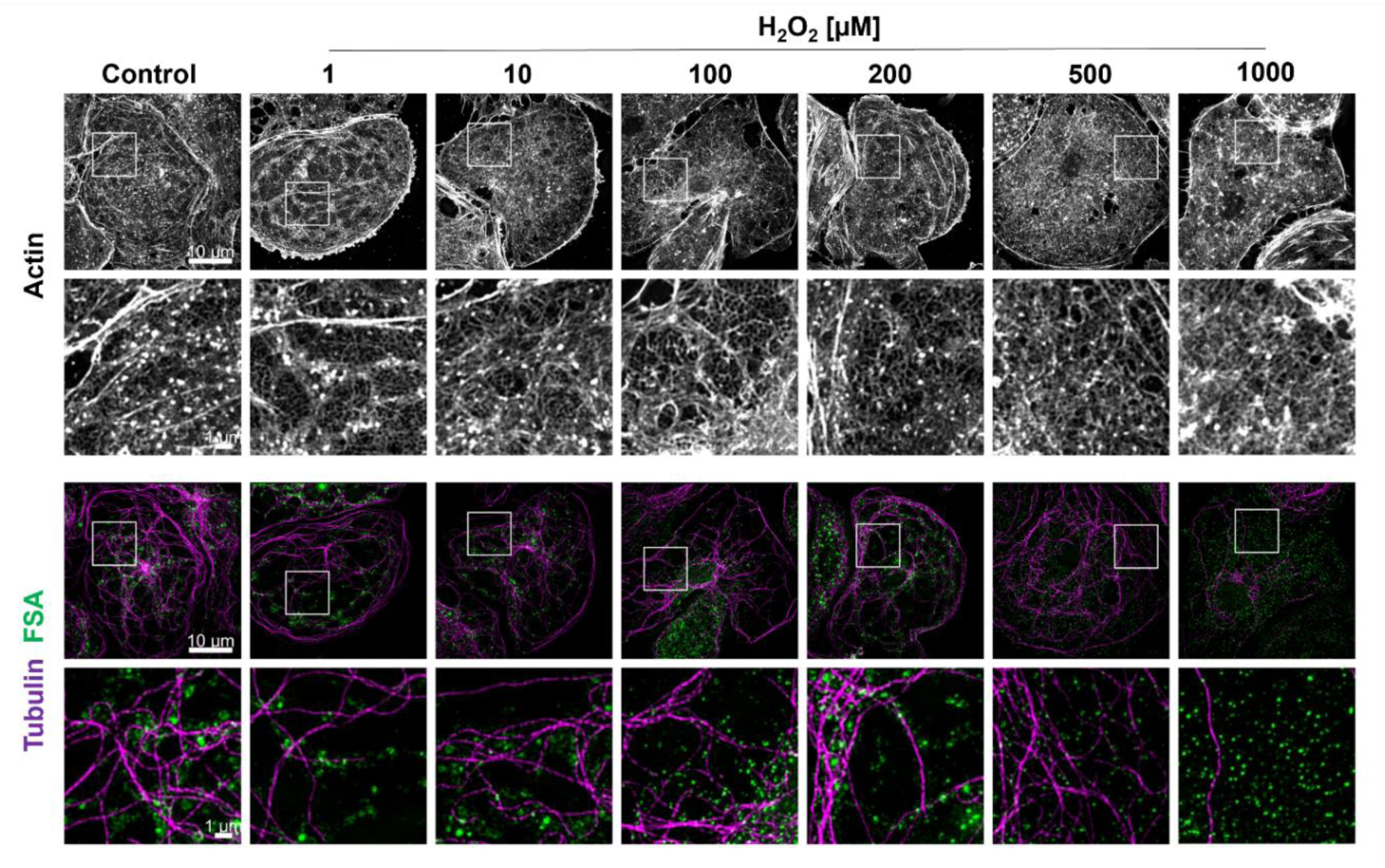
Representative SIM projection images of actin and tubulin cytoskeleton in rat LSEC treated with hydrogen peroxide. Cells were treated for 2 h with H_2_O_2_ with FSA-AF488 added for the last 15 min and after fixation stained with phalloidin-AF555 and anti-α-tubulin-AF647 antibody. The presented cells were selected based on the positive FSA signal indicating that the cell could still perform endocytosis. For H_2_O_2_ treatment above 50 μM only a fraction of LSEC population still can take up FSA as shown in overview images in Figure 1D. Bottom rows show high magnification images (10 x 10 µm) of the areas indicated in the top rows low magnification images (40.96 µm × 40.96 µm).

### ROS-depletion system in LSEC

To better understand the LSEC defence mechanisms against ROS, the effects of ROS-depleting agents were studied. A co- and pre-treatment with glutathione (GSH, 500µM) and n-acetyl cysteine (NAC, 0.5 and 2 mg/ml) together with 200 µM H_2_O_2_ were used and LSEC viability, internal ROS levels and scavenging activity were assessed (Figure 6). The concentration of hydrogen peroxide for these experiments was selected based on the results from the previous section (Figure 1-5) and showed a clear time-dependent reduction in functional and structural viability, a nearly 2-fold increase in intracellular ROS and reduced endocytic activity with completely inhibited FSA degradation.

**Figure 6.**
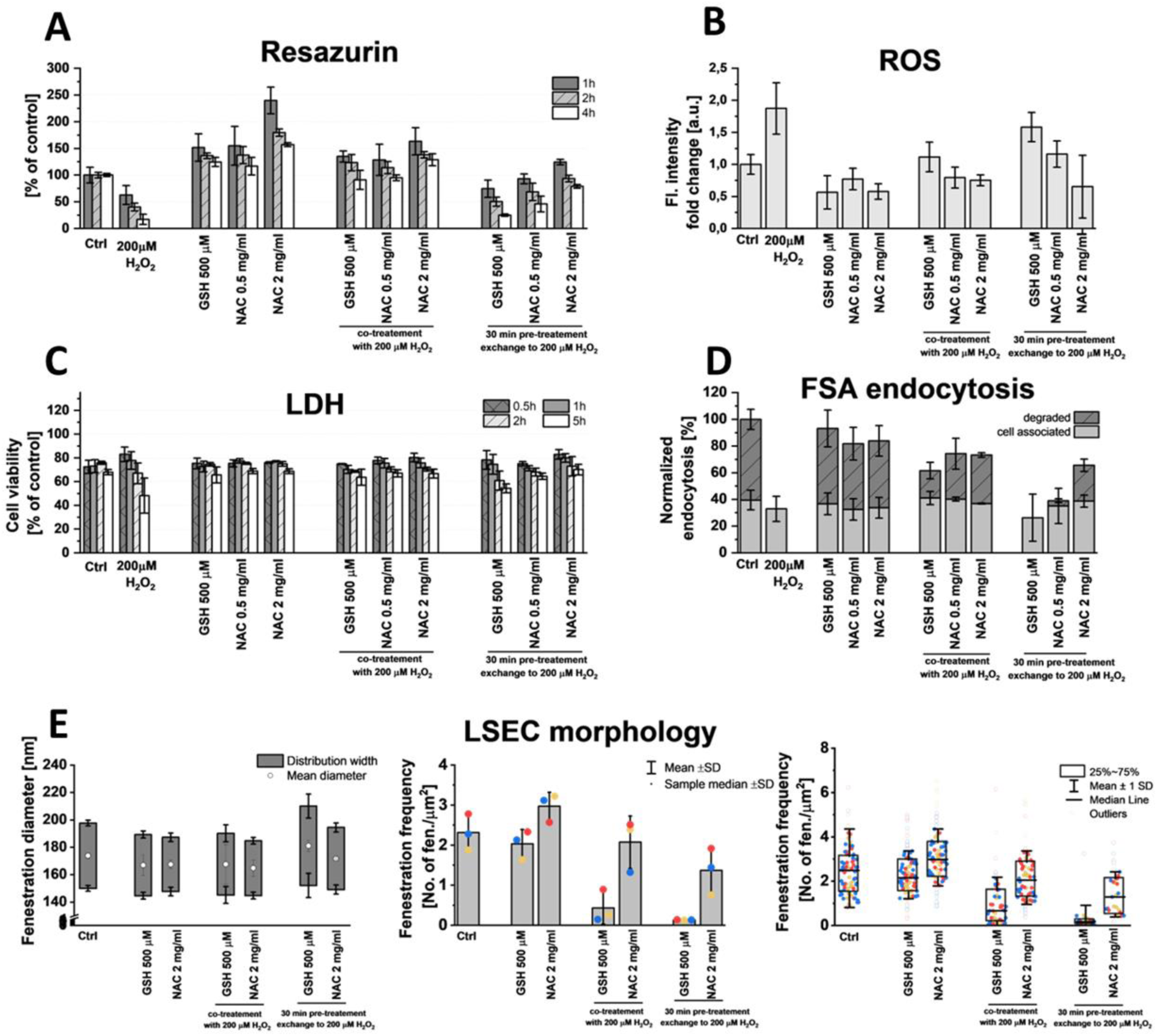
The effect of ROS-inducing vs. ROS-depleting agents on rat LSEC viability, functions and morphology. Cells were treated with ROS-inducing agent, hydrogen peroxide, or ROS-depleting agents: glutathione (GSH) or n-acetyl cysteine (NAC). For co-treatments, samples were simultaneously treated with both hydrogen peroxide and GSH/NAC at the starting point of the assays. For pre-treatments, samples were treated with GSH/NAC for 30 min, then rinsed with fresh media and treated with H_2_O_2_ at the starting point of the assays. The functional (A) and structural (C) viability were studied using resazurin and LDH release assays, respectively; (B) intracellular ROS levels were assessed using fluorescence-based ROS detection assay after 1h of treatment and (D) endocytic activity was measured with radiolabeled FSA-based scavenging assay during 2h treatment. (E) LSEC morphology was quantitatively assessed using SEM images. Colours represent data from separate bioreplicates.

All selected ROS-depleting agents were tested independently and showed no effect on structural viability (in LDH release assay) or endocytic activity (Figure 6C/D). GSH and NAC decreased the intracellular ROS and significantly increased the reduction potential of the cells as shown in the functional viability resazurin assay (statistical results can be found in Table *S5*, compared to control and Table S7, time-dependent, in the supplemental information). The resazurin assay measures mitochondrial activity as a reduction potential of the cell, which as expected is increased when exposed to ROS-depleting agents such as GSH and NAC which provide cells with additional reducing power. LSEC morphology was not affected by GSH, while NAC treatment led to a 30% increase in the number of fenestrations without changes in fenestration diameters (Figure 6E).

In comparison with the hydrogen peroxide challenge alone, simultaneous treatment with GSH or NAC showed a similar reduction of negative ROS effects. Co-treatment with NAC (both 0.5 and 2 mg/ml) almost completely mitigates the decrease in viability and intracellular ROS production and keeps the endocytic activity above 80% of control. GSH co-treatment, although reducing the effect of H_2_O_2_, still led to an increase in intracellular ROS and a decrease in endocytic activity, especially degradation. The cell associated fractions did not show any significant differences in comparison to the control. For the degraded fractions, treatment with 200 µM H_2_O_2_ together with all pre-treatments, and the GSH co-treatment showed a significant difference to control, indicating no reduction in the effect of H_2_O_2_ (Figure 6D; Table S8).

To exclude the effects from any direct interaction between GSH/NAC and hydrogen peroxide, sequential treatment was also used. In the samples with anti ROS pre-treatment, only the higher concentration of NAC prevented negative ROS effects, while lower concentration of NAC showed a reduction in intracellular ROS and LDH release but still led to a decrease in endocytic activity. Pre-treatment with GSH had no ROS-reducing effect and after following treatment with hydrogen peroxide an increase in intracellular ROS and decrease in viability and endocytic activity was observed (Figure 6B/C/D).

Based on the pre- and co-treatment viability and endocytosis results, only the higher, 2 mg/ml, concentration of NAC was used to study LSEC morphology (Figure 6E). The fenestration diameter distributions were not affected by the co-treatments with GSH/NAC and H_2_O_2_. 2h treatment with 200 µM H_2_O_2_ led to complete disruption of the cell membrane with no cells remaining for the morphological analysis of fenestrations (Figure 3). In the samples pre-treated with GSH, but not NAC, an increase in mean fenestration diameter and in fenestration diameter distribution width was observed. The fenestration frequency in samples with NAC co-treatment (2.1±0.5) remained on the same level as in control (2.3±0.3), while in pre-treatment samples a slight decrease (1.4±0.5) occurred. In cells co-treated with GSH, significant defenestration was observed, however, in comparison with treatment with hydrogen peroxide alone, the cell membranes remained intact in the majority of the cells. Pre-treatment with GSH led to significant, nearly complete loss of fenestrations, fenestration enlargement and gaps formation (see Table S9 for statistical analysis).

## Discussion

Cellular oxidative stress is defined as the imbalance between the production of ROS and reduction by various antioxidants. Elevated ROS correlates with metabolic syndrome [30], where ROS levels trend upwards with elevated BMI, and are reduced with weight loss [31]. A high fat/western diet causes an increase in ROS levels in both rat and mouse models [32][33]. In the liver, excessive ROS formation can occur in states of inflammation [34], by activated Kupffer cells and hepatic stellate cells, or in hepatocytes during intoxication events. LSEC were not reported to contribute to liver ROS production but due to their placement, LSEC can be exposed to high oxidative stress from both exogenous oxidants in the portal vein and other hepatic cells. Hydrogen peroxide is widely used to induce ROS formation in studying oxidative stress [35][36][37]. Physiologically, hydrogen peroxide is a product of mitochondrial metabolism and various H_2_O_2_ secretion levels were reported in the livers of different species [38][39].

In this study, we used H_2_O_2_ to generate intracellular ROS formation in LSEC *in vitro* and observed a clear pattern in the effects on cell viability, scavenging function and morphology for a wide range of hydrogen peroxide concentrations. The intraculluar ROS levels slightly increased in concentration of 5-100 µM but in a non-dose-dependent manner. Only for high concentrations of H_2_O_2_ above 100 µM a dose dependent increase in ROS was detected.

### Scavenging

LSEC are the main component of the body’s scavenger system, removing from the circulation grams amount of waste macromolecules per day [40][11][41]. Our results show that ROS can irreversibly reduce the endocytic activity even after very short exposure times. The cell’s degrading ability is affected first while with increasing concentrations also the cell-associated fraction decreases. Redox homeostasis has been shown to regulate lysosomal function [42] and increased ROS can hamper lysosomal function directly by preventing acidification and destabilizing lysosomal structure or indirectly by interfering with transport of endosomes. The tubulin network creates a highway for the transportation of endocytosed ligands to the lysosomes in LSECs (Falkowska-Hansen et al., 2007) and we observed deteriorated tubulin cytoskeleton, however, only for high concentrations of H_2_O_2_. For concentrations of 50-200 μM, we showed that the fraction of LSEC population that no longer can efficiently perform endocytosis is increasing rather than endocytic activity per cell is decreasing. These results suggest that for low concentrations of ROS the lysosomal function is primarily affected while with the increasing dose amount of ROS disruption of tubulin contributes to the reduction of scavenging function.

The impaired clearance by the scavenging system was found in ageing [43][44], as well as other liver diseases [40] and could be in part due to sustained inflammation and ROS generation by immune cells such as Kupffer cells [13]. This impairment of waste clearance is linked with liver disease related kidney injury [45], and could present an approach to prevention or amelioration via anti-oxidants. The interconnected nature of scavenging cells causes failures in one, especially LSEC, to cascade over to other systems - splenic and liver clearance of dead cells, cell remnants, bacteria and senescent erythrocytes depends on the same receptor system [46][47][48], thus impairments to the liver would impact spleen and bone marrow uptake as well by reducing capacity of the overall system [49]. Our data indicates NAC as an opportune candidate for cases with abnormally high oxidative stress.

### Morphology

We found that *in vitro* exposure to hydrogen peroxide reduces fenestration number in LSEC in a dose dependent manner during first 0.5 h of exposure, but the fenestrations reopen after about 1h before the membrane disintegrates towards 2 h or treatment. These findings corelate with the reports of Cogger et al. where rat livers were perfused with 70 and 700 µM H_2_O_2_ [50]. In that study, a decrease in the number of fenestration and thickening of endothelium was observed after 10 min similarly to our results from live imaging with AFM (Figure 4). Moreover, the defenestration centres we observed destabilize LSEC structure and make them more prone to damage, which may explain the gap formation in the liver perfusion model of Cogger et al. In the report of Martinez et al., generation of endogenous H_2_O_2_ was related to faster rate of defenestration of rat LSEC in culture. LSEC were cultured in 20% vs 5% O_2_ and increased levels of H_2_O_2_ were measured after 24 h and 48 h in the high oxygen conditions which correlated with lower porosity [51]. LSEC defenestration and gap formation was also observed *in vivo* in mice with elevated oxidative stress associated with a high fat western diet [33].

Moreover, in live LSEC imaging under the influence of H_2_O_2_ we observed closing of fenestrations without disrupting the underlying fenestration associated cytoskeleton and loss of the dynamics. We previously showed similar effects in LSEC challenged with antimycin A [52] – a mitochondrial cytochrome c reductase inhibitor known for increasing ROS production [53], and with diamide [54] – a known cytoskeletal drug that disrupts spectrin [55]. The complete structure of LSEC fenestrations is not yet fully described, but these results suggest that the ROS induced defenestration is related to the oxidation and destabilization of spectrin, and possibly other proteins that connect the cell membrane to fenestra associated cytoskeleton. Protein disulfide isomerase A1 (PDIA1) was recently identified to regulate fenestration dynamics and PDIA1 inhibition led to significant reduction of fenestration number independently of the cytoskeleton [56]. Although, LSEC fenestrated morphology is related to the actin cytoskeleton [12], we also observed no effect of H_2_O_2_ on cytoskeleton indicating that the H_2_O_2_ effect is independent from actin.

### Viability

After exposure to hydrogen peroxide, LSEC first lose their reducing equivalents before cell death as shown by resazurin assay and LDH release, respectively. The effect is both dose and time dependent, with cell death showing clear threshold effects. The reducing equivalents are depleted first, comparing the same timepoints and concentrations, before the cells proceed towards cell death suggesting that LSEC survival depends on reducing equivalents to counteract the ROS. Similar observations have been made for ROS-mediated conditions such as sinusoidal obstruction syndrome [57][58] or DILI [59]. Intriguingly, the viability and reducing potential of LSEC when compared with morphology, indicate that the cells close their fenestrations as reducing equivalents are being consumed. This can potentially be a hepatoprotective mechanism against the sudden increase in ROS-generating factors in the environment, especially considering the first pass effect that exposes LSEC to higher than systemic plasma levels concentrations of potentially harmful substances absorbed via the gastrointestinal tract. The reopening of fenestration could delay the exposure of hepatocytes until the dilution of the stressors within systemic circulation. Nevertheless, more research is needed to explain this phenomenon as well as confirm it *in vivo*.

### NAC/GSH

A glutathione-based defense system in LSEC has been previously described to play a protective role in ischemia-reperfusion injury (IRI), virus infections, and drug induced liver toxicity [15]. In our study, we observed that the effects of hydrogen peroxide induced ROS can be mitigated by simultaneous treatment with GSH or GSH precursor – NAC. The results with pretreatments, where only NAC but not GSH reduced the negative H_2_O_2_ effects, suggest that LSEC do not store excess amount of GSH but rather can readily produce it in the occurrence of oxidative stress conditions in provided with the fuel such as e.g., NAC. Similar data linking depletion of endogenous GSH with exacerbated cytotoxicity and the addition of exogenous GSH with reduced toxicity were noted by Deleve et al. [60]. Similarly, in chemically induced sinusoidal obstruction syndrome models, ROS-related damage of LSEC can be prevented by cotreatment with GSH [58]. This suggests that the cells can survive for as long as they have reducing equivalents to counteract the ROS. The expenditure of reducing equivalents thus eventually leads to cell damage and death, if more than what cell can produce/regenerate. The intracellular GSH levels in LSEC are much lower in comparison with hepatocytes, 0.5-1.5 fmol/cell and 17-50 fmol/cell respectively [60][15], making LSEC more sensitive to oxidative stress. The limited reducing capability of LSEC can explain the observed in our study non-dose-dependent increase in intracellular ROS levels for lower H_2_O_2_ concentration of 5-100 µM, which suggests that the cell can successfully neutralise ROS until some threshold level. Only for high concentrations of H_2_O_2_ above 100 µM a dose-dependent increase in ROS was detected.

NAC is typically used as a hepatoprotective agent to prevent e.g., hepatic ischemia-reperfusion injury (HIRI)[61] or drug-induced liver injury (DILI)[62], especially acetaminophen overdose [63]. In both HIRI and DILI prevention, the reduction of oxidative stress and ROS production was shown to play a crucial role. Moreover, ROS are required for KC proinflammatory/antigen presenting activity [64] and NAC as well as other antioxidants decrease LPS-induced Kupffer cell activation and TNF-alpha secretion [65][66]. Our results with NAC pretreatment being protective against ROS suggest that the reduced LSEC toxicity is crucial to reducing overall hepatic toxicity in both HIRI and DILI. As shown by Cogger et al., acute exogenous oxidative stress leads to gap formation in LSEC and disruption of the endothelial layer, leading to further exposure of hepatocytes to the portal blood. The reduction of ROS-related toxicity in LSEC can help avoid further exposure of hepatocytes and alterations of the space of Disse preventing the toxicity for the whole organ.

### Conclusions

Hydrogen peroxide induced intracellular ROS formation and was used to study oxidative stress effects on rat liver sinusoidal endothelial cells (LSEC) *in vitro*. ROS irreversibly reduced LSEC endocytic/scavenging function, potentially due to disrupted tubulin cytoskeleton at high ROS levels. ROS caused a dose-dependent reduction in LSEC fenestrations within 0.5 h, followed by reopening before membrane disintegration around 2h. ROS-induced LSEC defenestration is possibly related to oxidation and destabilization fenestration ultrastructure, but independent of actin cytoskeleton. LSEC first lose reducing equivalents before undergoing cell death upon H_2_O_2_ exposure, suggesting fenestration closure as a potential hepatoprotective mechanism against sudden ROS increases also suggesting antioxidants as a potential therapeutic approach. NAC and GSH mitigate H_2_O_2_-induced ROS effects in LSEC, with NAC pretreatment being more effective, indicating LSEC can readily produce GSH under oxidative stress if provided with precursors such as NAC. NAC’s protective effects against ROS-mediated LSEC toxicity could contribute to its hepatoprotective role in conditions like ischemia-reperfusion injury and drug-induced liver injury, by preventing further hepatocyte exposure and alterations in the space of Disse.

## Acknowledgements

The authors would like to thank Randi Olsen and Tom-Ivar Eilertsen from Advanced Microscopy Core Facility at UiT for electron microscopy expertise and Professor Peter McCourt for linguistic revision of the manuscript.

This study was supported by EU project DeLIVERy EIC-2021-Pathfinder grant no. 101046928 and Hop-On Facility HORIZON-WIDERA program associated with DeLIVERy.

## Authors contributions: (https://credit.niso.org)

Conceptualization KS, CFH, LDK; Data curation CFH, LDK, KS, WH, BZ, ES; Formal analysis ES, LDK, CFH; Funding acquisition TH, BZ; Investigation CFH, LDK, KS, JS, WH, BZ; Methodology CFH, LDK, KS, JS, WH, BZ, ES; Supervision KS; Visualization CFH, LDK, KS, WH, BZ; Writing - original draft: CFH, LDK, KS; and Writing - review & editing: all authors.

## Data statement

The datasets used and/or analyzed during the current study and not provided in the manuscript/supplementary information are available from the corresponding author on reasonable request.

## Supplementary Informations

**Table S1.** Statistical Matrix with anova analysis for Figure 1A. Using Shapiro Wilks test the sample distribution of all conditions was defined as normally distributed. ANOVA for mean values was applied to assess the effect of H2O2 [µM] on the internal ROS formation.

| Treatment (vs Control)<br>[H<sub>2</sub>O<sub>2</sub> [μM]] | p.value | significance |
| --- | --- | --- |
| 1 | 0.81 | ns |
| 10 | 0.084 | ns |
| 20 | 0.0028 | ** |
| 50 | 0.048 | * |
| 100 | 8.22E-06 | *** |
| 200 | 7.30E-13 | *** |
| 500 | 5.31E-14 | *** |
| 1000 | 2.53E-14 | *** |

**Table S2.1.** P for trend analysis for Error! Reference source not found.B. Using Shapiro Wilks test the sample distribution of all conditions was defined as normally distributed. Linear model for p for trend was applied over mean values for different H2O2 concentrations ranging from 0 to 1000 µM. Conditions were split based on observable trend difference between 0-50 and 50-1000 µM when the full model (0-1000µM) was significant.

Cell associated
| Treatment<br>H<sub>2</sub>O<sub>2</sub> [μM] | Slope | R<sup>2</sup> | p for trend | Significance |
| --- | --- | --- | --- | --- |
| All | -0.04 | 0.43 | 1.00009E-09 | *** |
| 0-50 | -0.11 | 0.025 | 0.31 | ns |
| 50-1000 | -0.025 | 0.32 | 0.00058 | *** |
| Degraded |  |  |  |  |
| Treatment<br>H<sub>2</sub>O<sub>2</sub> [μM] | Slope | R<sup>2</sup> | p for trend | Significance |
| All | -0.046 | 0.37 | 2.57E-08 | *** |
| 0-50 | -0.65 | 0.43 | 2.59E-06 | *** |
| 50-1000 | -0.013 | 0.20 | 0.0090 | ** |

**Table S2.2.**
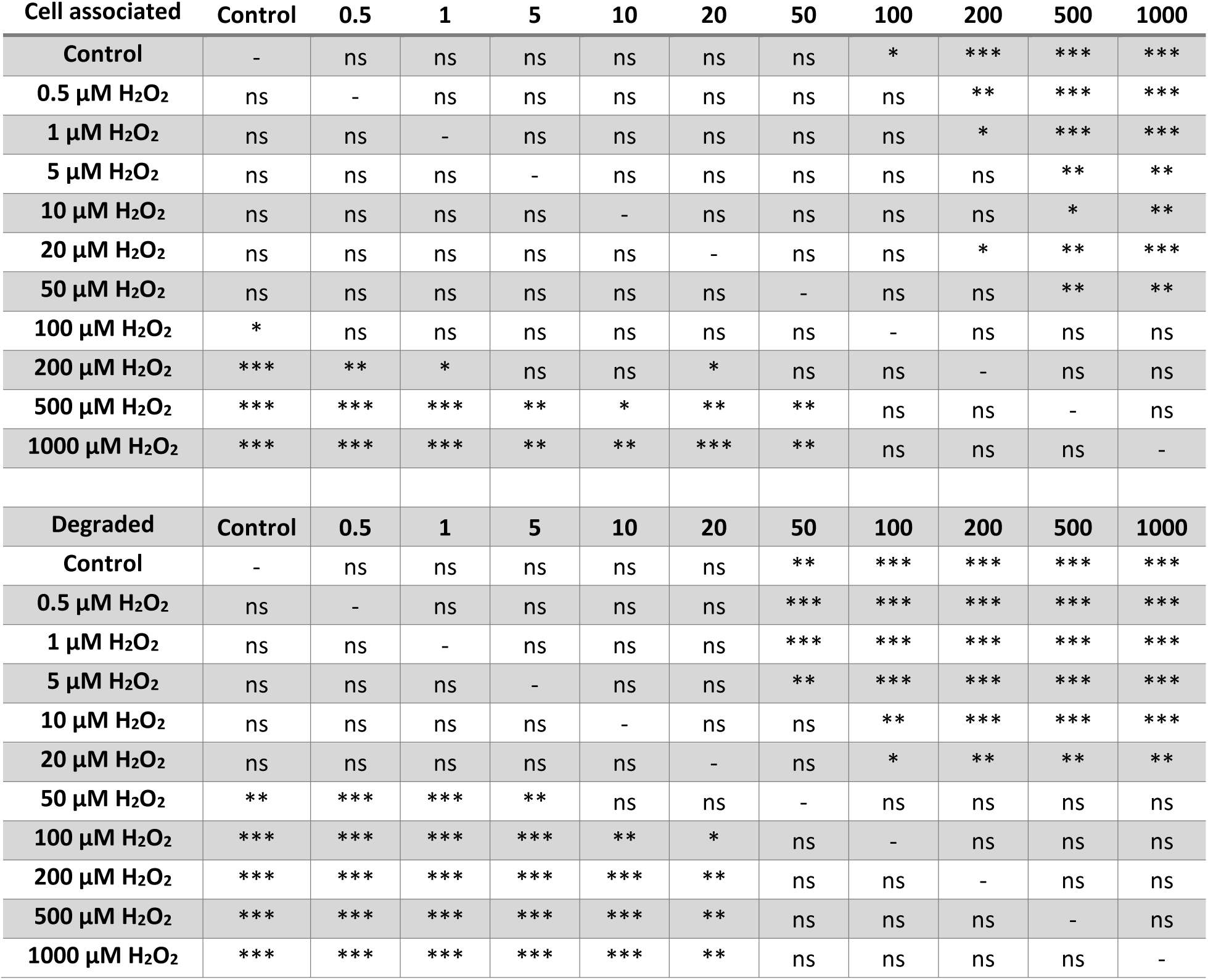
Statistical Matrix with anova analysis for Error! Reference source not found.B. Using Shapiro Wilks test the sample distribution of all conditions was defined as normally distributed. ANOVA for mean values was applied to assess the effect of H_2_O_2_ [µM] treatment on FSA endocytosis, divided into cell associated and degraded fractions.

| Cell associated | Control | 0.5 | 1 | 5 | 10 | 20 | 50 | 100 | 200 | 500 | 1000 |
| --- | --- | --- | --- | --- | --- | --- | --- | --- | --- | --- | --- |
| Control | - | ns | ns | ns | ns | ns | ns | * | *** | *** | *** |
| 0.5 μM H<sub>2</sub>O<sub>2</sub> | ns | - | ns | ns | ns | ns | ns | ns | ** | *** | *** |
| 1 μM H<sub>2</sub>O<sub>2</sub> | ns | ns | - | ns | ns | ns | ns | ns | * | *** | *** |
| 5 μM H<sub>2</sub>O<sub>2</sub> | ns | ns | ns | - | ns | ns | ns | ns | ns | ** | ** |
| 10 μM H<sub>2</sub>O<sub>2</sub> | ns | ns | ns | ns | - | ns | ns | ns | ns | * | ** |
| 20 μM H<sub>2</sub>O<sub>2</sub> | ns | ns | ns | ns | ns | - | ns | ns | * | ** | *** |
| 50 μM H<sub>2</sub>O<sub>2</sub> | ns | ns | ns | ns | ns | ns | - | ns | ns | ** | ** |
| 100 μM H<sub>2</sub>O<sub>2</sub> | * | ns | ns | ns | ns | ns | ns | - | ns | ns | ns |
| 200 μM H<sub>2</sub>O<sub>2</sub> | *** | ** | * | ns | ns | * | ns | ns | - | ns | ns |
| 500 μM H<sub>2</sub>O<sub>2</sub> | *** | *** | *** | ** | * | ** | ** | ns | ns | - | ns |
| 1000 μM H<sub>2</sub>O<sub>2</sub> | *** | *** | *** | ** | ** | *** | ** | ns | ns | ns | - |
| Degraded | Control | 0.5 | 1 | 5 | 10 | 20 | 50 | 100 | 200 | 500 | 1000 |
| Control | - | ns | ns | ns | ns | ns | ** | *** | *** | *** | *** |
| 0.5 μM H<sub>2</sub>O<sub>2</sub> | ns | - | ns | ns | ns | ns | *** | *** | *** | *** | *** |
| 1 μM H<sub>2</sub>O<sub>2</sub> | ns | ns | - | ns | ns | ns | *** | *** | *** | *** | *** |
| 5 μM H<sub>2</sub>O<sub>2</sub> | ns | ns | ns | - | ns | ns | ** | *** | *** | *** | *** |
| 10 μM H<sub>2</sub>O<sub>2</sub> | ns | ns | ns | ns | - | ns | ns | ** | *** | *** | *** |
| 20 μM H<sub>2</sub>O<sub>2</sub> | ns | ns | ns | ns | ns | - | ns | * | ** | ** | ** |
| 50 μM H<sub>2</sub>O<sub>2</sub> | ** | *** | *** | ** | ns | ns | - | ns | ns | ns | ns |
| 100 μM H<sub>2</sub>O<sub>2</sub> | *** | *** | *** | *** | ** | * | ns | - | ns | ns | ns |
| 200 μM H<sub>2</sub>O<sub>2</sub> | *** | *** | *** | *** | *** | ** | ns | ns | - | ns | ns |
| 500 μM H<sub>2</sub>O<sub>2</sub> | *** | *** | *** | *** | *** | ** | ns | ns | ns | - | ns |
| 1000 μM H<sub>2</sub>O<sub>2</sub> | *** | *** | *** | *** | *** | ** | ns | ns | ns | ns | - |

**Table S3.** P for trend analysis for Error! Reference source not found.C. Using Shapiro Wilks test the sample distribution of all conditions was defined as normally distributed. Linear model for p for trend was applied over mean values for different H_2_O_2_ concentrations ranging from 0 to 500 µM for different times points: 10 min / 30 min treatment with H_2_O_2_ and 30 min treatment with a recovery time of 2h /10h and split by fraction (degraded and cell associated). Time: treatment times with H2O2; recovery: time to recover from treatment with H_2_O_2_.

| Fraction | Time [min] | Recovery [h] | Intercept | Slope | R<sup>2</sup> | p value | Significance |
| --- | --- | --- | --- | --- | --- | --- | --- |
| Degraded | 10 | 0 | 54.10 | -20.61 | 0.98 | 2.01E-07 | *** |
| Cell Associated | 10 | 0 | 45.89 | 5.45 | 0.85 | 0.051 | ns |
| Degraded | 30 | 0 | 53.32 | -14.027 | 0.98 | 1.58E-05 | *** |
| Cell Associated | 30 | 0 | 46.67 | 11.31 | 0.92 | 0.00027 | *** |
| Degraded | 30 | 2 | 59.08 | -23.93 | 0.98 | 4.15E-07 | *** |
| Cell Associated | 30 | 2 | 40.91 | 3.35 | 0.89 | 0.33 | ns |
| Degraded | 30 | 10 | 59.48 | -23.17 | 0.99 | 1.03E-07 | *** |
| **Cell Associated** | 30 | 10 | 40.51 | -2.09 | 0.96 | 0.46 | ns |

**Table S4.** Anova significance testing for Error! Reference source not found.H/I. Only biologically significant pairs are listed here. All other combinations are not significantly different. Time 1 and treatment 1 are one analysis point, compared to time 2 and treatment 2.

| Time1 [h] | Treatment1<br>[μM H<sub>2</sub>O<sub>2</sub>] | Time2 [h] | Treatment2<br>[μM H<sub>2</sub>O<sub>2</sub>] | p.adj | Significance |
| --- | --- | --- | --- | --- | --- |
| 1 | 500 | 1 | 0 | 0.029 | * |
| 1 | 500 | 0.5 | 500 | 0.00021 | *** |

**Table S5.** P for trend analysis for Error! Reference source not found.H/I. Using Shapiro Wilks test the sample distribution of all conditions was defined as normally distributed. Linear model for p for trend was applied over mean values for fenestration frequency (number of fenestrations per µm^2^) at different H_2_O_2_ concentrations (ranging from 0 to 500 µM) for three treatment lengths, 30 min, 1h and 2h.

| Time [h] | Slope | R<sup>2</sup> | P value | Significance |
| --- | --- | --- | --- | --- |
| 0.5 | -0.0035 | 0.52 | 0.00021 | *** |
| 1 | -0.0036 | 0.59 | 4.59e-05 | *** |
| 2 | -0.0037 | 0.43 | 0.0013 | ** |

**Table S6.** Anova significance testing for Error! Reference source not found.A. Comparison between the control and different treatments for different time points (1h, 2h, 4h). NAC05: NAC 0.5mg/mL; NAC2: 2mg/mL. Pretreatment: 30 min pre treated cells with NAC/GSH, then H_2_O_2_ was added. Co-treatment: GSH/NAC and H_2_O_2_ were added at the same time.

| Treatment A | Time A | Treatment B | Time B | p.value | Significance |
| --- | --- | --- | --- | --- | --- |
| Control | 1 | NAC 05; 200μM H<sub>2</sub>O<sub>2</sub> cotreatment | 1 | 0.43 | ns |
| Control | 1 | NAC 05; 200μM H<sub>2</sub>O<sub>2</sub> pretreatment | 1 | 1 | ns |
| Control | 1 | NAC 2; 200μM H<sub>2</sub>O<sub>2</sub> pretreatment | 1 | 0.76 | ns |
| Control | 1 | NAC05 | 1 | 1.94e-05 | *** |
| Control | 1 | NAC2 | 1 | 0 | *** |
| Control | 1 | NAC2; 200μM H<sub>2</sub>O<sub>2</sub> cotreatment | 1 | 2.13e-07 | *** |
| Control | 1 | 200 μM H<sub>2</sub>O<sub>2</sub> | 1 | 0.042 | * |
| Control | 1 | 500μM GSH | 1 | 9.19e-05 | *** |
| Control | 1 | 500μM GSH; 200μM H<sub>2</sub>O<sub>2</sub> cotreatment | 1 | 0.091 | ns |
| Control | 1 | 500μM GSH; 200μM H<sub>2</sub>O<sub>2</sub> pretreatment | 1 | 0.69 | ns |
| Control | 2 | NAC 05; 200μM H<sub>2</sub>O<sub>2</sub> cotreatment | 2 | 0.99 | ns |
| Control | 2 | NAC 05; 200μM H<sub>2</sub>O<sub>2</sub> pretreatment | 2 | 0.22 | ns |
| Control | 2 | NAC 2; 200μM H<sub>2</sub>O<sub>2</sub> pretreatment | 2 | 1 | ns |
| Control | 2 | NAC05 | 2 | 0.041 | * |
| Control | 2 | NAC2 | 2 | 1.23e-11 | *** |
| Control | 2 | NAC2; 200μM H<sub>2</sub>O<sub>2</sub> cotreatment | 2 | 0.031 | * |
| Control | 2 | 200 μM H<sub>2</sub>O<sub>2</sub> | 2 | 1.49e-06 | *** |
| Control | 2 | 500μM GSH | 2 | 0.060 | ns |
| Control | 2 | 500μM GSH; 200μM H<sub>2</sub>O<sub>2</sub> cotreatment | 2 | 0.84 | ns |
| Control | 2 | 500μM GSH; 200μM H<sub>2</sub>O<sub>2</sub> pretreatment | 2 | 0.00025 | *** |
| Control | 4 | NAC 05; 200µM H<sub>2</sub>O<sub>2</sub> cotreatment | 4 | 1 | ns |
| **Control** | **4** | **NAC 05; 200µM H<sub>2</sub>O<sub>2</sub> pretreatment** | **4** | **2.82e-05** | **\*\*\*** |
| Control | 4 | NAC 2; 200µM H<sub>2</sub>O<sub>2</sub> pretreatment | 4 | 0.94 | ns |
| Control | 4 | NAC05 | 4 | 0.99 | ns |
| **Control** | **4** | **NAC2** | **4** | **6.10e-06** | **\*\*\*** |
| Control | 4 | NAC2; 200µM H<sub>2</sub>O<sub>2</sub> cotreatment | 4 | 0.41 | ns |
| **Control** | **4** | **200 µM H<sub>2</sub>O<sub>2</sub>** | **4** | **1.91e-12** | **\*\*\*** |
| Control | 4 | **500µM GSH** | 4 | 0.76 | ns |
| Control | 4 | 500µM GSH; 200µM H<sub>2</sub>O<sub>2</sub> cotreatment | 4 | 0.99 | ns |
| **Control** | **4** | **500µM GSH; 200µM H<sub>2</sub>O<sub>2</sub> pretreatment** | **4** | **2.20e-10** | **\*\*\*** |

**Table S7.** Anova significance testing for Error! Reference source not found.A. Comparison in between the same treatment for different time points (1h, 2h, 4h). Treatment 1 and treatment 1 are one analysis point, compared to time 2/treatment 2. NAC05: NAC 0.5mg/mL; NAC2: 2mg/mL. Pretreatment: 30 min pre treated cells with NAC/GSH, then H_2_O_2_ was added. Co-treatment: GSH/NAC and H_2_O_2_ were added at the same time.

| Treatment A | Time A [h] | Treatment B | Time B [h] | p.value | significance |
| --- | --- | --- | --- | --- | --- |
| 200 µM H<sub>2</sub>O<sub>2</sub> | 1 | 200 µM H<sub>2</sub>O<sub>2</sub> | 2 | 0.88 | ns |
| 200 µM H<sub>2</sub>O<sub>2</sub> | 1 | 200 µM H<sub>2</sub>O<sub>2</sub> | 4 | 0.0016 | ** |
| 200 µM H<sub>2</sub>O<sub>2</sub> | 2 | 200 µM H<sub>2</sub>O<sub>2</sub> | 4 | 0.85 | ns |
| 500µM GSH | 1 | 500µM GSH | 2 | 0.99 | ns |
| 500µM GSH | 1 | 500µM GSH | 4 | 0.54 | ns |
| 500µM GSH | 2 | 500µM GSH | 4 | 0.99 | ns |
| 500µM GSH; 200µM H<sub>2</sub>O<sub>2</sub> cotreatment | 1 | 500µM GSH; 200µM H<sub>2</sub>O<sub>2</sub> cotreatment | 2 | 0.99 | ns |
| 500µM GSH; 200µM H<sub>2</sub>O<sub>2</sub> cotreatment | 1 | 500µM GSH; 200µM H<sub>2</sub>O<sub>2</sub> cotreatment | 4 | 0.0034 | ** |
| 500µM GSH; 200µM H<sub>2</sub>O<sub>2</sub> cotreatment | 2 | 500µM GSH; 200µM H<sub>2</sub>O<sub>2</sub> cotreatment | 4 | 0.19 | ns |
| 500µM GSH; 200µM H<sub>2</sub>O<sub>2</sub> pretreatment | 1 | 500µM GSH; 200µM H<sub>2</sub>O<sub>2</sub> pretreatment | 2 | 0.78 | ns |
| 500µM GSH; 200µM H<sub>2</sub>O<sub>2</sub> pretreatment | 1 | 500µM GSH; 200µM H<sub>2</sub>O<sub>2</sub> pretreatment | 4 | 0.00026 | *** |
| 500µM GSH; 200µM H<sub>2</sub>O<sub>2</sub> pretreatment | 2 | 500µM GSH; 200µM H<sub>2</sub>O<sub>2</sub> pretreatment | 4 | 0.70 | ns |
| NAC 05; 200µM H<sub>2</sub>O<sub>2</sub> cotreatment | 1 | NAC 05; 200µM H<sub>2</sub>O<sub>2</sub> cotreatment | 2 | 0.99 | ns |
| NAC 05; 200µM H<sub>2</sub>O<sub>2</sub> cotreatment | 1 | NAC 05; 200µM H<sub>2</sub>O<sub>2</sub> cotreatment | 4 | 0.14 | ns |
| NAC 05; 200µM H<sub>2</sub>O<sub>2</sub> cotreatment | 2 | NAC 05; 200µM H<sub>2</sub>O<sub>2</sub> cotreatment | 4 | 0.98 | ns |
| NAC 05; 200µM H<sub>2</sub>O<sub>2</sub> pretreatment | 1 | NAC 05; 200µM H<sub>2</sub>O<sub>2</sub> pretreatment | 2 | 0.74 | ns |
| NAC 05; 200µM H<sub>2</sub>O<sub>2</sub> pretreatment | 1 | NAC 05; 200µM H<sub>2</sub>O<sub>2</sub> pretreatment | 4 | 0.00075 | *** |
| NAC 05; 200µM H<sub>2</sub>O<sub>2</sub> pretreatment | 2 | NAC 05; 200µM H<sub>2</sub>O<sub>2</sub> pretreatment | 4 | 0.88 | ns |
| NAC 2; 200µM H<sub>2</sub>O<sub>2</sub> pretreatment | 1 | NAC 2; 200µM H<sub>2</sub>O<sub>2</sub> pretreatment | 2 | 0.27 | ns |
| NAC 2; 200µM H<sub>2</sub>O<sub>2</sub> pretreatment | 1 | NAC 2; 200µM H<sub>2</sub>O<sub>2</sub> pretreatment | 4 | 0.0016 | ** |
| NAC 2; 200µM H<sub>2</sub>O<sub>2</sub> pretreatment | 2 | NAC 2; 200µM H<sub>2</sub>O<sub>2</sub> pretreatment | 4 | 0.99 | ns |
| NAC05 | 1 | NAC05 | 2 | 0.99 | ns |
| NAC05 | 1 | NAC05 | 4 | 0.031 | * |
| NAC05 | 2 | NAC05 | 4 | 0.95 | ns |
| NAC2 | 1 | NAC2 | 2 | 1.22e-06 | *** |
| NAC2 | 1 | NAC2 | 4 | 2.09e-12 | *** |
| NAC2 | 2 | NAC2 | 4 | 0.88 | ns |
| NAC2; 200µM H<sub>2</sub>O<sub>2</sub> cotreatment | 1 | NAC2; 200µM H<sub>2</sub>O<sub>2</sub> cotreatment | 2 | 0.71 | ns |
| NAC2; 200µM H<sub>2</sub>O<sub>2</sub> cotreatment | 1 | NAC2; 200µM H<sub>2</sub>O<sub>2</sub> cotreatment | 4 | 0.10 | ns |
| NAC2; 200µM H<sub>2</sub>O<sub>2</sub> cotreatment | 2 | NAC2; 200µM H<sub>2</sub>O<sub>2</sub> cotreatment | 4 | 0.99 | ns |
| Control | 1 | Control | 2 | 1 | ns |
| Control | 1 | Control | 4 | 1 | ns |
| Control | 2 | Control | 4 | 1 | ns |

**Table S8.** Anova significance testing for Error! Reference source not found.D. Comparison of endocytosis of Iodine 125-coupled FSA between control (RPMI cultured cells) and different (co/pre) treatments. Separated into cell associated and degraded fractions. NAC05: NAC 0.5mg/mL; NAC2: 2mg/mL. Pretreatment: 30 min pre-treated cells with NAC/GSH, then H_2_O_2_ was added. Co-treatment: GSH/NAC and H_2_O_2_ were added at the same time.

| Treatment A | Treatment B | p.value | Significance |
| --- | --- | --- | --- |
| **Cell Associated** |  |  |  |
| Control | NAC 05 | 0.95 | ns |
| Control | NAC 05; 200µM H<sub>2</sub>O<sub>2</sub> cotreatment | 0.99 | ns |
| Control | NAC 05; 200µM H<sub>2</sub>O<sub>2</sub> pretreatment | 0.99 | ns |
| Control | NAC 2 | 0.97 | ns |
| Control | NAC2; 200µM H<sub>2</sub>O<sub>2</sub> cotreatment | 0.99 | ns |
| Control | NAC2; 200µM H<sub>2</sub>O<sub>2</sub> pretreatment | 0.99 | ns |
| Control | 200µM H<sub>2</sub>O<sub>2</sub> | 0.96 | ns |
| Control | 500µM GSH | 0.99 | ns |
| Control | 500µM GSH; 200µM H<sub>2</sub>O<sub>2</sub> cotreatment | 0.99 | ns |
| Control | 500µM GSH; 200µM H<sub>2</sub>O<sub>2</sub> pretreatment | 0.62 | ns |
| **Degraded** |  |  |  |
| Control | NAC 05 | 0.99 | ns |
| Control | NAC 05; 200µM H<sub>2</sub>O<sub>2</sub> cotreatment | 0.19 | ns |
| Control | NAC 05; 200µM H<sub>2</sub>O<sub>2</sub> pretreatment | 5.93e-05 | *** |
| Control | NAC 2 | 0.99 | ns |
| Control | NAC2; 200µM H<sub>2</sub>O<sub>2</sub> cotreatment | 0.27 | ns |
| Control | NAC2; 200μM H<sub>2</sub>O<sub>2</sub> pretreatment | 0.035 | * |
| Control | 200μM H<sub>2</sub>O<sub>2</sub> | 3.47e-05 | *** |
| Control | 500μM GSH | 0.99 | ns |
| Control | 500μM GSH; 200μM H<sub>2</sub>O<sub>2</sub> cotreatment | 0.0061 | ** |
| Control | 500μM GSH; 200μM H<sub>2</sub>O<sub>2</sub> pretreatment | 4.28e-05 | *** |

**Table S9.** Anova significance testing for Error! Reference source not found.E – Fenestration frequency. Comparison between the control and different treatments. NAC05: NAC 0.5mg/mL; NAC2: 2mg/mL. Pretreatment: 30 min pre-treated cells with NAC/GSH, then H_2_O_2_ was added. Co-treatment: GSH/NAC and H_2_O_2_ were added at the same time.

| Treatment A | Treatment B | p.value | significance |
| --- | --- | --- | --- |
| Control | 2 mg/mL NAC | 0.85 | ns |
| Control | 500μM GSH | 0.86 | ns |
| Control | NAC2; 200μM H<sub>2</sub>O<sub>2</sub> cotreatment | 0.79 | ns |
| Control | NAC2; 200μM H<sub>2</sub>O<sub>2</sub> pretreatment | 0.075 | ns |
| Control | 500μM GSH; 200μM H<sub>2</sub>O<sub>2</sub> cotreatment | 0.0055 | ** |
| Control | 500μM GSH; 200μM H<sub>2</sub>O<sub>2</sub> pretreatment | 0.00016 | *** |

**Figure S1.**
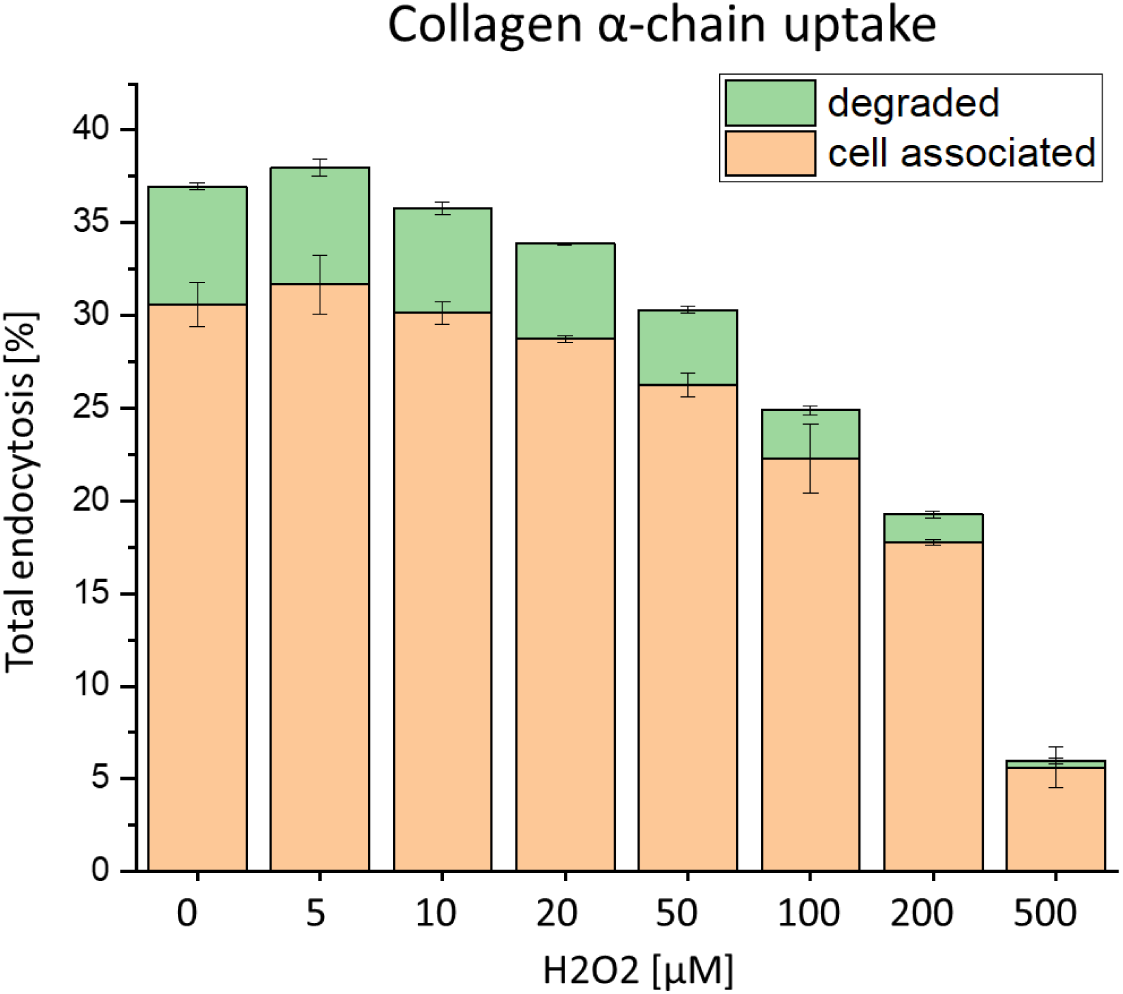
Effects of ROS on endocytosis of trace amount of ^125^I-alpha-chain collagen in rat liver sinusoidal endothelial cell (LSEC). Cells were treated with hydrogen peroxide together radiolabeled collagen for 2h in RPMI. Green bars represent degraded ligand while orange bars represent cell-associated fraction of the ligand (±SD), n=2 bio replicates.

**Figure S2.**
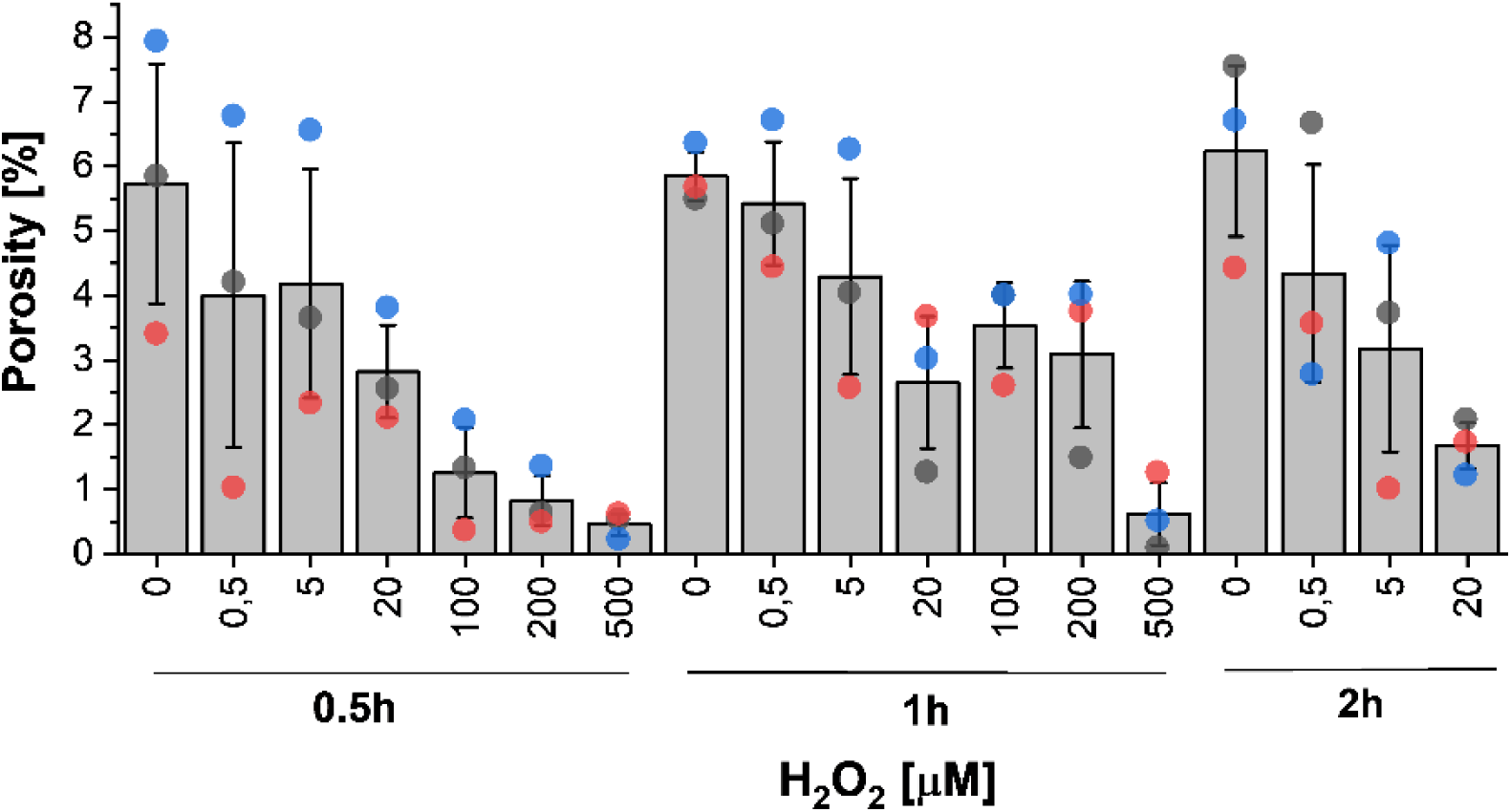
Changes in LSEC porosity after exposure to H2O2. Each dot represents the mean porosity value calculated from each bioreplicate (n=3), bar correspond to mean of means ± SD.

